# Learning-induced neurobiological mechanisms enabled by RGFP966 enhance signal-in-noise responsivity in auditory cortex and behavior

**DOI:** 10.64898/2025.12.15.694497

**Authors:** Nilay Ateşyakar, Kasia M. Bieszczad

**Author notes:** **Corresponding author**: Dr. Kasia M. Bieszczad, 327 Psychology Building, Rutgers University-New Brunswick 152 Frelinghuysen Road Piscataway, New Jersey 08854 USA.

## Abstract

Auditory learning enables sound-selective enhancements in auditory cortical (AC) processing. Background noise can also alter sound-selective auditory responsivity. Yet, how learning can enhance AC processing in noise is unknown. Pharmacological inhibition of histone deacetylase 3 (HDAC3) via RGFP966 enhances learning and related AC plasticity, but its potential to support signal detection under degraded acoustic conditions is unclear. To determine if task learning supports tone-signal detection in a later background noise challenge, adult rats (Sprague-Dawley males) were trained in ideal quiet conditions to learn a tone-reward association while treated with RGFP966 (TRAINED+RGFP966, n=6). RGFP966 accelerated sound-reward learning relative to untreated rats (TRAINED, n=5), though all animals reached equivalent high levels of performance before further testing. Successful performance produced *sound-specific* enhancements in AC responses evoked by the learned tone, and a *sound-general* effect that suppressed responses to noise, relative to untrained rats (NAÏVE, n=7). Notably, frequency-selective response biases were latent under quiet conditions and became robustly expressed under background noise, particularly in RGFP966-treated learners who acquired the task more rapidly. Increasing background noise abolished frequency-selective enhancements in tone-evoked AC activity, yet the learning-induced suppressive effect to noise was maintained. Behavioral detection of the learned tone across noise conditions mirrored AC tone-evoked response patterns. The findings demonstrate that learning can engage coordinated cortical mechanisms regulated by HDAC3 that selectively modify representations of behaviorally relevant signals. Further, auditory memory is dynamically gated by sensory context, relying on the stability of cortical decoding mechanisms to support listening in real-world environments.

**SIGNIFICANCE STATEMENT:** Difficulty hearing in noise is a widespread challenge, yet the cortical mechanisms that preserve meaningful sounds under noisy listening conditions remain unclear. We show that HDAC3 inhibition via the pharmacological inhibitor, RGFP966, accelerates auditory learning and strengthens cortical encoding of a learned tone while suppressing background noise activity in ways that predict improved behavioral detection. These findings reveal an experience-driven cortical mechanism that supports improved hearing in challenging listening environments, which may inform strategies for enhancing auditory learning and rehabilitation using HDAC3 drug-targets.

## INTRODUCTION

A major function of the auditory system is to identify behaviorally relevant acoustic signals in the environment. Successful sound-guided performance depends on neural mechanisms that enhance the fidelity and selectivity of sound representations beyond that explained by peripheral signal-to-noise ratio (SNR) alone. Associative learning drives central plasticity in the auditory cortex (AC), producing lasting modifications to receptive field tuning, excitatory–inhibitory circuit balance, and population-level coding (Weinberger, 2004; Schreiner and Polley, 2014; Froemke, 2015). Learning enhances encoding of behaviorally relevant sounds (Fritz et al., 2003) and contributes to the formation of precise auditory memories (Bieszczad and Weinberger, 2010; McGann, 2015). However, it remains unknown how learning-driven changes in cortical coding could support sound-guided behavior under degraded listening conditions such as background noise. Despite its relevance to real-world auditory function, the neural representation of learned sound signals in background noise remains poorly understood.

Background noise affects auditory discrimination (Alain et al., 2014). Noise increases the sound levels required for tone-evoked activity and decreases response magnitudes (Phillips, 1985; Phillips and Hall, 1986). Human word recognition declines with increasing noise even when SNR is held constant (Studebaker et al., 1999; Dubno et al., 2005). In animal models, noise reduces neural population response diversity (Shilling-Scrivo et al., 2021) and modulates tuning selectivity (Teschner et al., 2016). Yet in some cases, noise can enhance frequency discrimination by reducing overlap in cortical tuning (Christensen et al., 2019). Indeed, cortical inhibition plays a central role in modulating receptive fields (Lakunina et al., 2022). For example, activating parvalbumin-expressing interneurons can mimic noise-induced reductions in responsivity and enhance discrimination of spectrally similar tones (Christensen et al., 2019; Nocon et al., 2023). Inhibitory circuits also contribute to the formation and maintenance of learning-induced AC plasticity and to the consolidation of long-lasting auditory memories (Aizenberg et al., 2015; Blackwell and Geffen, 2017; Natan et al., 2017; Wood et al., 2017; Lakunina et al., 2022; Studer and Barkat, 2022). These findings support that changes in cortical receptive field activity regulates the system’s capacity to separate spectral signals from noise. Learning-induced receptive field plasticity in auditory cortex may therefore underlie signal-in-noise performance gains for behaviorally relevant sounds. Although short-term AC plasticity can adaptively shape frequency-specific responses during task performance, enduring improvements in signal processing require molecular mechanisms that reliably stabilize experience-dependent changes over time.

Epigenetic modulation consolidates experience-dependent sensory cortical plasticity by driving enduring changes to cue-selective evoked activity and supporting persistent cue memories. Among the epigenetic regulatory framework, histone deacetylase 3 (HDAC3) has emerged as a particularly robust regulator of auditory cortical plasticity, with direct consequences for the selectivity and robustness of sound signal processing (Bieszczad et al., 2015). For example, HDAC3 inhibition during auditory associative learning enhances frequency-specific tuning and expands tonotopic regions that represent a learned signal frequency, relative to identically trained untreated animals (Shang et al., 2019; Rotondo and Bieszczad, 2020, 2021a, 2021b). Remarkably, enhancements in frequency-specific AC selectivity mediated by HDAC3-inhibition corresponds with sharply frequency-specific behavioral responding that persists for weeks after training. Treatment likely promotes long-lasting effects of learning via transcriptional control of cortical function (Graham et al., 2023). While HDAC3 inhibition reliably increases sound-specific behavior in silent backgrounds, its ability to maintain behavioral performance in novel noisy environments remains unexplored. HDAC3 inhibition may create an *a priori* advantage for processing a learned sound signal and protect signal-specific AC responses in novel background noise. Accordingly, HDAC3 manipulation provides an opportunity to identify cortical coding features that optimize signal-in-noise detection.

The present study examines how memory for a learned sound signal is selectively expressed in auditory cortex and behavior in the presence of background noise, and whether HDAC3 inhibition modulates its expression. By characterizing how learning-induced cortical plasticity interacts with noise-dependent auditory processing, this work addresses neurobiological mechanisms that support sound-guided behavior under ecologically relevant listening conditions.

## MATERIALS AND METHODS

### Subjects

Adult male Sprague Dawley rats weighing 250-300 grams on arrival served as subjects in this experiment in 3 groups: (1) TRAINED, n=15; (2) TRAINED+RGFP966, n=10; and (3) NAÏVE, n=7. Animals that were to undergo training were water-restricted and maintained at 85% of the free-drinking weight of littermate controls for the duration of training and memory testing. Untrained rats (the NAÏVE group) were never water restricted. Water supplementation was administered as needed to maintain weight and was given *ad libitum* prior to any surgical procedures. All rats were handled daily by an experimenter after arrival to the laboratory for at least 5 days before beginning any other procedures to acclimate animal subjects to transport to the lab, experimental handling, and the ambient sounds of the laboratory. The vivarium maintained a 12:12 light/dark cycle (lights on at 7am; lights off at 7pm). All procedures were approved and conducted in accordance with guidelines by the Institutional Animal Care and Use Committee at Rutgers, The State University of New Jersey (#PROTO999900026 to K.M.B.), in accordance with relevant named guidelines and regulations, and are reported in compliance with ARRIVE guidelines.

### Behavioral Training

Behavioral sessions were conducted in three identical instrumental conditioning chambers (Habitest Modular Test Cage System, 12”x10”x12” chamber; this and all Habitest modular components were from Coulbourn Instruments, Lehigh PA) fitted with a bar floor (H10-11R-TC-SF) within a custom-built sound-attenuating box. Each chamber was fitted with a response lever (H21-03R), house light (H11-01R), infrared lights (H27-91R), speaker (H12-01R) and a water delivery system (H14-05R) along the same wall. When triggered either by the response lever or a hand switch (H21-01), the water delivery system would present a water cup (0.1cc of water) in the reward port (1.26” W x 1.625”H). Daily training sessions were counterbalanced to ensure equal exposure to different chambers. All behavioral protocols were created in and run using Graphic State 4 software. Behavioral responses were recorded at a resolution of 1 millisecond. Auditory stimuli were generated using Tucker-Davis Technologies (TDT) hardware (RZ6) and RPvdsEx software (TDT; Alachua, FL) at a 50 kHz sampling rate and presented via the operant chamber’s wall-mounted speaker.

Rats were first behaviorally shaped to lever press for water rewards (1:1 fixed ratio) in daily 45-minute sessions across five consecutive days. Twenty-four hours following successful shaping in which animals independently initiated and maintained bar-press responding for water rewards across the 45-minute session or to satiation, animals began daily training sessions on a tone-reward associative learning task. During each 45-minute training session, tone trials were presented such that on each trial, animals heard a 5.0-kHz pure tone (60 dB SPL; 8 sec) and any lever presses made during the tone duration were reinforced with water reward (3 second access to the water delivery dipper system). Responses during silent inter-trial intervals (ITI; range: 5–25 seconds, randomized) resulted in an error signal (flashing house light) and a “time-out” a flashing house light error signal that initiated a 6-second time-out period that delayed the time until the next tone trial. Task acquisition was evaluated using two complementary performance criteria: First, an 80% performance criterion was defined as the first day on which the session performance (performance = [lever presses during tones / all bar presses during session] × 100) exceeded 80% and remained above 80% on the following day of tone-reward training. This metric was used to quantify learning speed. Second, asymptotic performance was defined using more stringent stability-based criteria to ensure high levels of stable performance prior to memory testing and electrophysiological recording: (1) a performance exceeding 90% on at least two of the last three training days within a maximum of 20 training days, and (2) performance stability, measured as a coefficient of variation below 10% across the final five training days. Animals proceeded to memory testing only after reaching asymptotic performance, or after >20 sessions of daily training. This performance criterion ensured stable, high-level task performance at the time of evaluating memory specificity and later electrophysiological recording in AC. Only 1 animal (from the TRAINED group) failed to achieve this performance criterion, so training was halted after 20 daily sessions of tone-reward task training.

### Pharmacological HDAC3 inhibition via RGFP966

To examine the role of HDAC3 inhibition in auditory learning, a subset of animals that comprised the TRAINED+RGFP966 group received systemic post-training injections of the selective HDAC3 inhibitor, RGFP966 (10 mg/kg, subcutaneous; ApexBio #A8803) immediately following training sessions on days 2, 3 and 4. Animals that were to undergo injections were exposed to the same handling procedure used to perform the injections for at least 5 days prior to injections (e.g., immediately following bar-press shaping sessions and after the first daily session (day 1) of tone-reward training) to reduce effects of novelty or arousal that later could have been induced by the injection itself. Trained animals that were not injected with RGFP966 (the TRAINED group) included animals that received either vehicle injections (equivalent volume, subcutaneous) immediately after training sessions on days 2, 3 and 4; or no injections at all across training. Vehicle injections did not produce differences in task acquisition in animals that were not injected (Wilcoxon rank-sum test: W=30.5, p=0.72; mean day to reach 80%; **Fig. S1**), so these animals were collapsed into a single (TRAINED) group. That injections themselves did not alter task performance is important to reinforce that any effects of injecting RGFP966 were due to the effects of the HDAC3 inhibitor itself, and not confounded by indirect effects of experiencing a subcutaneous injection.

### Memory Test

Upon meeting the performance criteria, a behavioral memory test was conducted to assess individual differences in noise-sensitive and frequency-selective behavior. To ensure stable task performance immediately prior to the Memory Test, animals completed a brief version of the tone–reward task, which lasted approximately 5 minutes (mean = 4 min 52 sec ± 29 sec). This brief session also served as a warm-up that was immediately followed by the Memory Test. Each animal received 117 pseudorandomized trials (three blocks of 39) in which the signal tone was presented (8 sec) at either the signal frequency (5.0 kHz) or ±¼ octave away at two nearby novel frequencies (4.2 and 5.9 kHz). Intertrial intervals were variable (5–20 sec). Tone presentations during the Memory Test were not reinforced with water rewards. Tones were delivered either in quiet or embedded in notched white noise at three levels that produced a defined signal-to-noise ratio: low noise (60 dB tone + 20 dB noise; SNR40), medium noise (60 dB tone + 40 dB noise; SNR20), and high noise (60 dB tone + 60 dB noise; SNR0, see **Fig. S2**). Noise was Gaussian white noise regenerated from a normal distribution on every trial (frequency range <25kHz; gated with 20 ms cosine rise/fall ramps). The notch filter was introduced at the frequency of the tone embedded in noise on every single trial to ensure that animals could not use an increase in spectral energy with frequency channels to guide their sound-cued behavior. As such, bar-presses during tone presentations were interpreted to be driven by activating an internal representation of the learned association between a specific tone frequency and the availability of reward.

Noise-sensitive and frequency-selective behavioral performance was analyzed separately for each noise condition within individual subjects before determining group-level effects. For each noise level, frequency-selective responding was quantified as the percentage of bar presses made during tone presentations to a given frequency relative to the total number of bar presses made during tone presentations of all frequencies tested within the same noise condition. This was, computed as: Proportion of Bar Presses % = (total # of bar presses made during tone A / total # of bar presses made during any tone at that SNR) × 100.

Finally, to ensure that the unrewarded context of memory testing did not catastrophically disrupt performance prior to electrophysiological recording, all animals went through a final task session of the tone-reward learning paradigm 24-48 hours later to confirm maintained task performance. All animals maintained performance from the last training session prior to memory testing (t(10) = 0.92, p = 0.37; paired samples t-test; **Fig. S3A**). Further, the final performance level measured after memory testing did not differ between groups (t(9) = 1.13, p = 0.28; independent samples t-test; **Fig. S3B**), which indicated that RGFP966 did not produce any effects on the maintenance of task performance following the memory testing session.

### Surgical Procedures

Following behavioral testing, *in vivo* multiunit electrophysiological recordings targeting auditory cortex (AC) were performed. All surgical procedures were conducted in a custom designed double-walled, walk-in sound-attenuated chamber (Industrial Acoustics Co., Bronx, NY). Animals were anesthetized with sodium pentobarbital (60 mg/kg, i.p.), with supplemental doses (10-15 mg/kg, i.p.) administered as needed to maintain a surgical plane of anesthesia. Atropine methyl nitrate (10 mg/kg, i.m.) was administered to minimize bronchial secretions. Ophthalmic ointment (Puralube®) was applied to prevent corneal desiccation. Core body temperature was maintained near 36–37°C throughout the procedure using a heating pad closed-loop homoeothermic warming system, continuously monitored via a rectal temperature probe (TC-1000, CWE Inc.). While under anesthesia, the head was shaved to reveal the scalp and the animal was mounted in a modified stereotaxic frame that tilted to provide access to the temporal plane (Kopf Instruments, Tujunga, CA). To provide local anesthesia, 2% lidocaine (LidoJect; Henry Schein; ∼1 mL, s.c.) was administered in small boluses across the scalp. The scalp was then resected, and the skull was secured to a head holder using several small stainless-steel screws and dental acrylic. This fixation allowed removal of the ear bars while maintaining the head in a stable and fixed position for the duration of the procedure. Animals were then secured in a stereotaxic apparatus. The right temporal muscle was retracted to expose the skull overlying auditory cortex. A craniotomy was performed over the right temporal cortex, and the dura mater was carefully removed to expose the cortical surface, which was maintained using warm sterile 0.9% saline application throughout the cortical mapping procedure to prevent cortical desiccation. The cisterna magna was periodically drained of cerebrospinal fluid to reduce cerebral swelling and improve cortical stability as needed.

### Auditory Cortical Mapping Procedures

To characterize the frequency-selective responsiveness of auditory cortical sites within the tonotopic organization primary auditory cortex (A1), animals underwent a terminal auditory cortical mapping procedure that followed established protocols described previously (Bieszczad & Weinberger, 2010a, 2010b, 2012; Bieszczad et al., 2015; Elias et al., 2015; Shang et al., 2019). Extracellular responses to acoustic stimuli were recorded across auditory cortex using parylene-coated tungsten microelectrode arrays consisting of six (2 × 3) channels (FHC, Bowdoin, ME; 1–2 MΩ impedance; 250 µm inter-electrode spacing). Electrodes were advanced orthogonal to the cortical surface into the thalamorecipient middle cortical layers (III–IV; approximately 400–600 µm depth) using a precision microdrive (IVM-1000; Scientifica, UK). Multiple penetrations (M=85.66 sites, SE=10.41 per animal in Phase 1; M=32.33 sites, SE=3.94 per animal in Phase 2; see Phase definitions in *Electrophysiological Recording Procedures*) were distributed across the cortical surface to obtain comprehensive spatial sampling of tone-responsive sites across the auditory cortex (AC). Photographs of the cortical surface were acquired prior to each penetration (OCS-SK2-14X; OptixCam Summit K2) to document electrode placement. Images were subsequently aligned using vascular landmarks to reconstruct a relative spatial map of recording sites and to delineate the borders within AC tonotopic organization.

### Electrophysiological Recording Procedures

All acoustic stimuli were delivered monaurally to the contralateral (left) ear via a single calibrated electromagnetic speaker (MF1; Tucker-Davis Technologies Inc.) positioned approximately 10 cm from the ear canal. Recordings at each cortical site were conducted in two sequential phases. Trial structure and stimulus timing were adapted from a previous study demonstrating the effects of background noise on auditory cortical frequency processing (Teschner et al., 2016).

**Phase 1:** In the first phase of the recording sequence, a search stimulus paradigm was used to confirm recording locations were bone fide sites within primary auditory cortex (A1) defined by short-latency responses to pure tones within the confines of the tonotopic organization of low to high frequencies from posterior to anterior locations across the temporal axis of the cortical plane. In this phase, tones were only presented in a quiet background to assess short-latency, well-tuned frequency-selective responses characteristic of A1. Pure tones spanned a 0.525–47.700 kHz frequency range (in quarter-octave spacing; 25 ms duration gated with 5 ms cosine rise/fall ramps), presented pseudorandomly at sound levels ranging from 10 to 60 dB SPL (in 10-dB steps), with 6 repetitions of each of the 162 frequency-level stimuli. This generated 972 number of total trials across the recording that included 27 frequencies and 6 sound levels. Inter-stimulus intervals were randomly jittered between 585-865 ms in 52 ms steps.

**Phase 2:** If the recording site could be confirmed as A1, the second phase of the recording sequence would proceed. In this phase, tone-evoked responses were recorded under different SNR conditions. Tones were identical to the first phase frequency range 0.525-47.700 kHz (quarter octave steps, 25 ms duration gated with 5 ms cosine ramps) at 10-60 dB SPL (10 dB steps). However in this recording phase, acoustic stimuli could be embedded in white Gaussian noise (i.e., regenerated from a Gaussian distribution on every trial; frequency range <25kHz) presented at three defined and fixed signal-to-noise ratios. Pure tones were presented either in quiet (SNR∞) or embedded in noise at fixed signal-to-noise ratios of low noise (SNR40) and high noise (SNR20). Each sound trial began with a pre-tone noise segment gated with 10 ms cosine ramps, followed by the onset of a 25 ms pure tone gated with 5 ms cosine rise/fall ramps that was effectively embedded within the noise duration. Noise continued for an additional 150 ms following tone offset. The onset latency of the tone relative to noise onset was at least 275 ms to generate a steady-state noise response, and varied pseudorandomly on different trials to be between 275 and 415 ms (in 20 ms increments). This generated a jitter so that tone onsets could not be predicted by the onset of the noise on each trial. Every trial concluded with a 275 ms silent interval to allow the system to return to an effective baseline, which ultimately yielded a fixed inter-noise interval of 725 ms from noise onset on one trial to the following trial noise onset. Each of the 162 frequency-level combination stimuli was pseudorandomly presented in 8 repetitions within each SNR condition for a total of 3888 trials across this phase of the recording session.

Neural signals were routed via a ZIF cable connecting the array adaptor mounted on the animal’s head to a 16-channel analog headstage and amplifier (PZ5 NeuroDigitizer Amplifier; Tucker-Davis Technologies) located outside of the recording chamber. Signals were amplified (×1000), digitized at 24.414 kHz using RZ6-A-P1 and RZ5D BioAmp processors (Tucker-Davis Technologies), and stored for subsequent offline spike detection and analysis using custom MATLAB© scripts. Data collection and online stimulus presentation were controlled using Synapse™ software (Tucker-Davis Technologies). Neural activity was monitored in real time using TDT Synapse™ and OpenEx software to initially determine tone and frequency selectivity, and these data were also stored for subsequent offline analysis.

Offline, unfiltered recordings were digitally band-pass filtered between 300 and 3000 Hz to isolate multiunit activity relative to timestamps that indicated noise and tone onsets across the continuous time series data. When present, which was rare, line noise and its harmonics (60 Hz and multiple) were identified on a per-channel basis and attenuated prior to spike detection. Multiunit discharges were characterized using previously reported temporal and amplitude criteria (Elias et al., 2015). Briefly, spike detection was performed using a window-discrimination method. For each recording site, detection thresholds were defined relative to the root mean square (RMS) of the signal computed from 500 randomly selected traces. Positive and negative detection thresholds were set at 1.5×RMS and 2.0×RMS, respectively. Accepted spike waveforms were required to exhibit peak-to-peak separations of no more than 0.6 ms. Spike waveforms associated with large-amplitude transients exceeding predefined amplitude thresholds were excluded to minimize contamination by electrical or unexpected and rare movement-related artifacts.

For each recording site, tone-evoked firing rates (spikes/s) were calculated by subtracting activity measured during a 40 ms window immediately prior to tone onset (−40 to 0 ms, with 0 defined as tone onset) from tone-evoked spiking measured during a 40 ms response-onset window (6–46 ms following tone onset). Responses exceeding ±1.0 SEM of the pre-tone firing rate were classified as evoked. Tone-evoked activity was used to construct frequency-response areas (FRAs) for each recording site, representing mean sound-evoked activity across all frequency-intensity combinations.

A behaviorally untrained and pharmacologically untreated control group (NAÏVE; n = 7) underwent identical electrophysiological recording procedures to establish baseline auditory cortical response properties in the absence of experimental training or pharmacological treatment beyond experiences of experimenter handling and housing within a standard vivarium and laboratory environment. These data were included in group comparisons that were ultimately conducted between three groups: NAÏVE, TRAINED, TRAINED+RGFP966.

### Frequency-selective AC Analyses

Analyses of frequency-selective effects in AC focused on activity evoked by the three behaviorally tested frequencies (4.2, 5.0, and 5.9 kHz) presented at either 60 or 40 dB SPL. To facilitate group-level comparisons across recording sites and experimental conditions, evoked firing rates were Z-scored within each signal-to-noise ratio condition. All analyses were performed using custom MATLAB© scripts. See *Results* for details.

### Code Accessibility

Custom Matlab© scripts used to analyze behavioral and neurophysiological data within will be provided within a reasonable timeframe upon reasonable request. Raw neurophysiological datasets will also be made available to the scientific community upon reasonable request via the *RUcore* at Rutgers University (rucore.libraries.rutgers.edu/research/).

### Statistical Analysis – Behavioral Responses

Days to reach the 80% performance criterion was compared between groups using Wilcoxon rank-sum tests, and the proportion of rapid learners (≥80% criterion reached within 10 days) was evaluated using Fisher’s Exact tests. Behavioral responses during the Memory Test were analyzed using Linear Mixed-Effects Models with fixed effects of Frequency, Group, and their interaction, with planned contrasts assessing frequency-specific biases. Behavioral engagement across noise conditions was analyzed using mixed-design ANOVA.

### Statistical Analysis – AC Responses

Electrophysiological responses to tones were analyzed using Linear Mixed-Effects Models conducted separately for each SNR condition, with fixed effects of Frequency and Group (TRAINED, TRAINED+RGFP966, NAÏVE) and their interaction. Post-hoc comparisons were corrected using Tukey or Bonferroni procedures. Steady-state noise-evoked activity, extracted from the pre-tone noise window, was analyzed using one-way ANOVA with Group as the independent factor, followed by Bonferroni-corrected pairwise tests where appropriate. All comparisons used two-tailed tests unless otherwise noted when motivated by *a priori* hypothesis of directional effects between groups or frequency factors. Alpha levels were α = 0.05. Statistical analyses were conducted using MATLAB© (v2021b), R Statistical Software (v4.4.0), and JASP (v0.18.3).

## RESULTS

### Auditory learning is accelerated with post-training HDAC3 inhibition

Performance on the tone-reward associative task was evaluated across the full training cohort (n = 25). All rats successfully learned to bar-press to the tone cue for reward and ultimately reached high levels of asymptotic performance (**Fig. 1A**). As shown in Figure 1A, RGFP966-treated rats (n = 10/25) displayed earlier and faster acquisition trajectories than identically trained but untreated controls (n = 15/25), with group means diverging primarily during the early training days. RGFP966-treated rats reached the 80% performance criterion more rapidly than untreated rats (9.75 vs. 11.8 days; **Fig. 1B, C**). Therefore, a 10-day threshold was used to define rapid learners, i.e., as animals reaching criterion in fewer than 10 training days. In the full cohort, 7 of 10 RGFP966-treated rats and only 5 of 15 untreated rats met this threshold. Although this difference in proportions was not statistically significant (p = 0.11, odds ratio = 0.21, one-tailed Fisher’s Exact test; **Fig. 1D**), the cumulative distribution of criterion attainment displayed a visually clear leftward shift in the RGFP966 group, consistent with an effect of HDAC3 inhibition to accelerate learning (**Fig. 1C**). Further neural analyses were limited to animals with successful electrophysiological cortical mapping that made data available across the targeted frequency representations in primary auditory cortex (see *Methods*; n = 13/25). Within this mapped subset, more RGFP966-treated rats (6 out of 7) than untreated rats (1 out of 6) reached criterion within 10 days (p = 0.025, odds ratio = 0.03, one-tailed Fisher’s Exact test), mirroring the observed effect of HDAC3 inhibition across the full training cohort to accelerate learning. To avoid performance-related confounds that could introduce variability in analyses to identify the neural substrates of HDAC3-inhibition, the slowest-learning RGFP966-treated animal and the fastest-learning untreated animal were both excluded as outliers. Following these exclusions, neurophysiological analyses on AC frequency coding were conducted on a trained sub-group treated with the HDAC3-inhibitor (TRAINED+RGFP966; n = 6) and compared to the identically trained, but untreated group (TRAINED; n = 5). By definition, these groups differed in rate of task acquisition. RGFP966-treated rats were faster to learn the tone-reward association (Wilcoxon rank-sum test: W < 0.001, p = 0.007; **Fig. 1E, F**).

**Figure 1.**
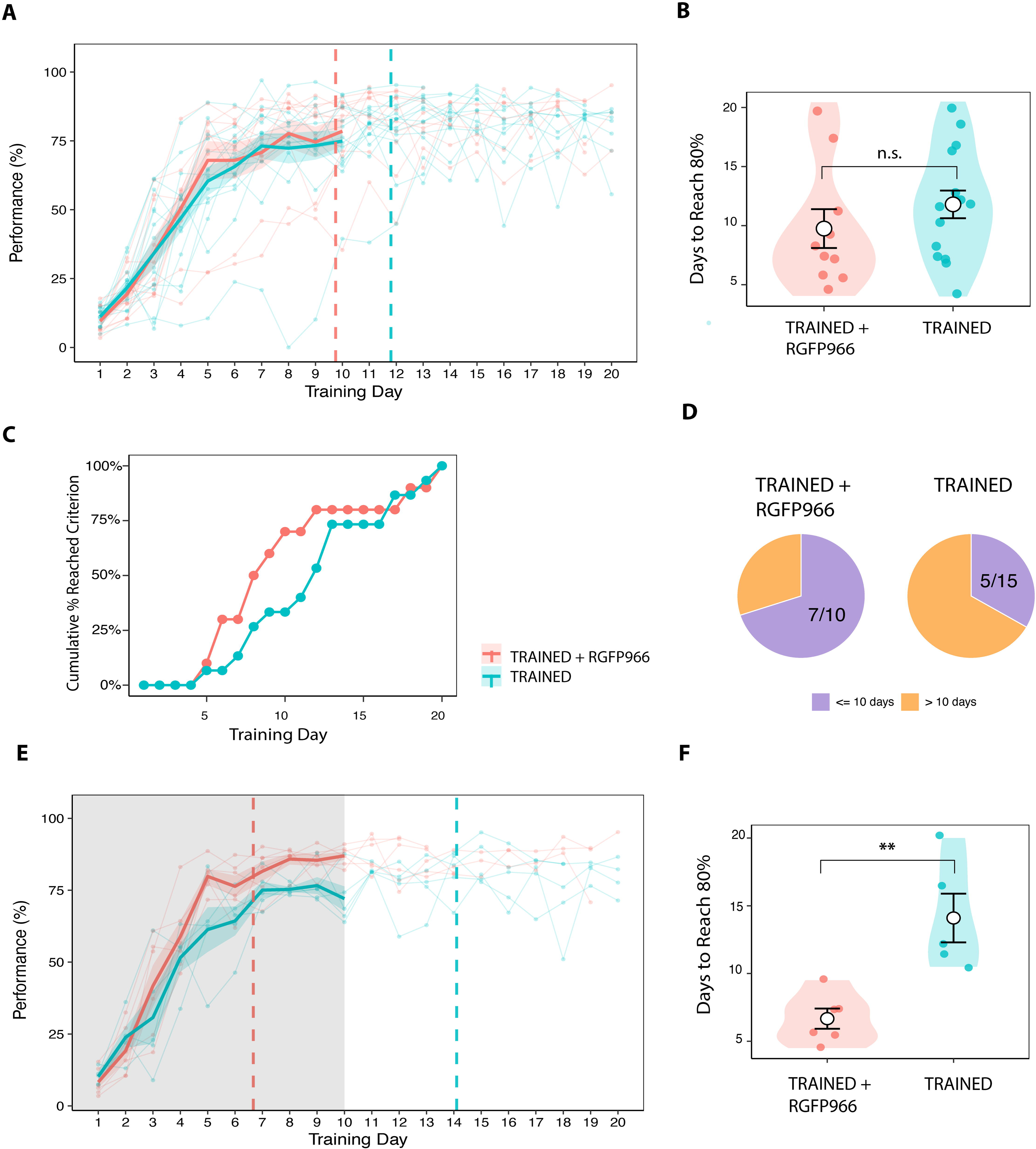
Auditory learning is accelerated by post-training HDAC3 inhibition. **(A)** Acquisition of tone–reward associative learning plotted as daily performance (% correct) across training days for the full cohort (n = 25). Thin lines represent individual animals; thick lines indicate group means (±SEM). Rats treated with the HDAC3 inhibitor RGFP966 (TRAINED+RGFP966; n = 10) exhibited earlier and steeper learning trajectories compared with identically trained but untreated controls (TRAINED; n = 15), with divergence most evident during early training. Dashed vertical lines indicate the mean day at which each group reached the 80% performance criterion. **(B)** Days required to reach the 80% performance criterion for RGFP966-treated and untreated rats in the full cohort. Individual animals are shown with group means ± SD overlaid; n.s., not significant. **(C)** Cumulative percentage of animals reaching the 80% criterion as a function of training day, demonstrating a leftward shift in the RGFP966-treated group indicative of accelerated learning. **(D)** Proportion of rapid learners, defined as animals reaching criterion in fewer than 10 training days, in the full cohort. Numbers within pie charts indicate animals reaching criterion in ≤10 days versus >10 days. **(E)** Acquisition curves for the subset of animals with successful auditory cortical mapping used for subsequent neurophysiological analyses (TRAINED+RGFP966, n = 6; TRAINED, n = 5), plotted as in A. Shaded region denotes the period used to assess acquisition rate, and dashed lines indicate mean criterion day for each group. **(F)** Days to reach the 80% criterion for the mapped subset shown in E. RGFP966-treated rats learned significantly faster than untreated rats (Wilcoxon rank-sum test; **p < 0.01).

### HDAC3 inhibition during training facilitates frequency-specific auditory cortical activity under low noise conditions

Frequency response areas (FRAs) were determined for each auditory cortical recording site in both trained groups and also in an untrained group of rats (NAÏVE; n = 7) to evaluate the effect of background noise on tone-evoked activity related to task learning. A region of interest within each FRA was defined as three test frequencies targeting the training tone frequency (i.e., 5.0kHz) and novel tones nearby (4.2kHz and 5.9kHz) either at the training tone sound level (i.e., n60 dB SPL) or at a lower sound level (40 dB SPL) for analysis in the three SNR conditions (**Fig. 2**). Spike rates were normalized (Z-scored sp/s) across recording sites to facilitate group-level analyses per SNR condition (SNR∞, SNR40, SNR20).

**Figure 2.**
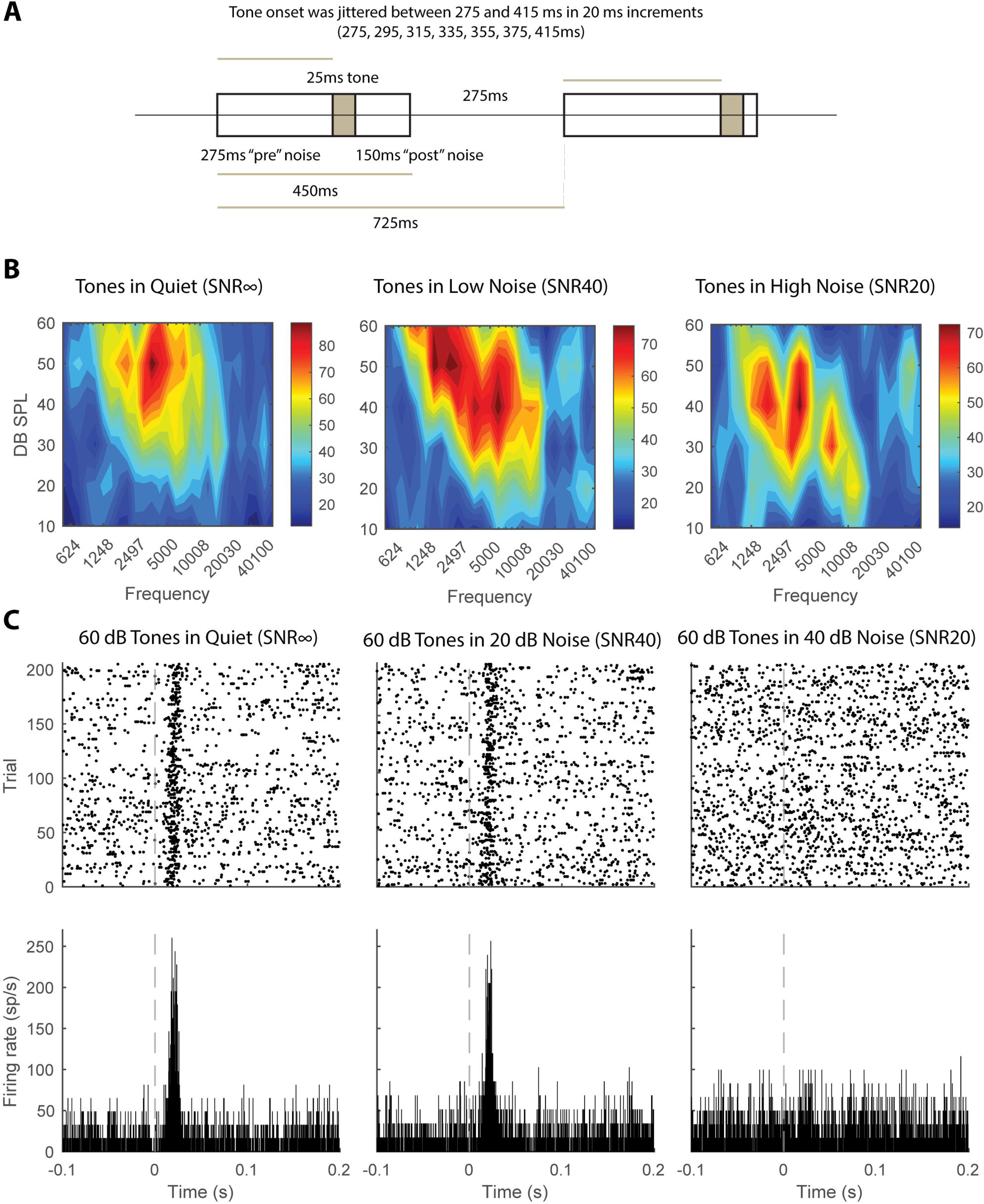
Definition of frequency response area (FRA) region of interest and effects of background noise on tone-evoked auditory cortical activity. **(A)** Schematic of the electrophysiological trial structure used to assess tone-evoked responses in auditory cortex. A 25-ms pure tone was presented on each trial, embedded in either quiet (SNR∞) or background noise (SNR20 or SNR40). Tone onset was temporally jittered between 275 and 415 ms (20-ms increments) relative to trial start to reduce anticipatory responses. Background noise preceded and followed tone presentation, yielding fixed pre- and post-tone noise epochs for response estimation. **(B)** Example frequency response areas (FRAs) from a representative auditory cortical recording site obtained under quiet (SNR∞), low-noise (SNR40), and high-noise (SNR20) conditions. Color maps depict tone-evoked firing rate (spikes/s) as a function of tone frequency and sound level (dB SPL). The black boxed region indicates the predefined region of interest (ROI) used for quantitative analyses at 60 dB SPL tones. **(C)** Corresponding spike rasters (top row) and peristimulus time histograms (PSTHs; bottom row) for responses to 60-dB SPL tones presented at the training frequency within each SNR condition. Dashed vertical lines indicate tone onset.

In quiet (SNR∞), the 5.9 kHz tone elicited the strongest relative evoked activity in all groups (60 dBs, Frequency: F(2,771) = 8.86, p = 0.00016; Group: F(2,771) < 0.001, p = 1.00; Frequency x Group: F(4,771) = 2.24, p = 0.062). Responses to 5.9 kHz were significantly higher than responses to 4.2 kHz (post hoc Linear Mixed-Effects Model contrasts collapsed across groups (Tukey-corrected) estimate = -0.298, t(514) = -4.16, p = 0.0001) and higher than responses to 5.0 kHz (estimate = -0.19, t(514) = -2.65, p = 0.022; **Fig. 3A** and **Table 1**). The AC response bias to 5.9 kHz was not evident when presented at sound levels below the 60 dB training sound level, e.g., at 40 dBs (40 dBs, Frequency: F(2, 771) = 0.14, p = 0.86). Similarly, just as there was no significant effect of training or of the HDAC3-inhibitor at the training tone sound level, there were no significant effects at the lower sound level (40dBs, Group: F(2, 771) < 0.001 p = 1.00; Group x Frequency: F(4, 771) = 0.24 p = 0.91; **Fig. 3B** and **Table 2)**.

**Figure 3.**
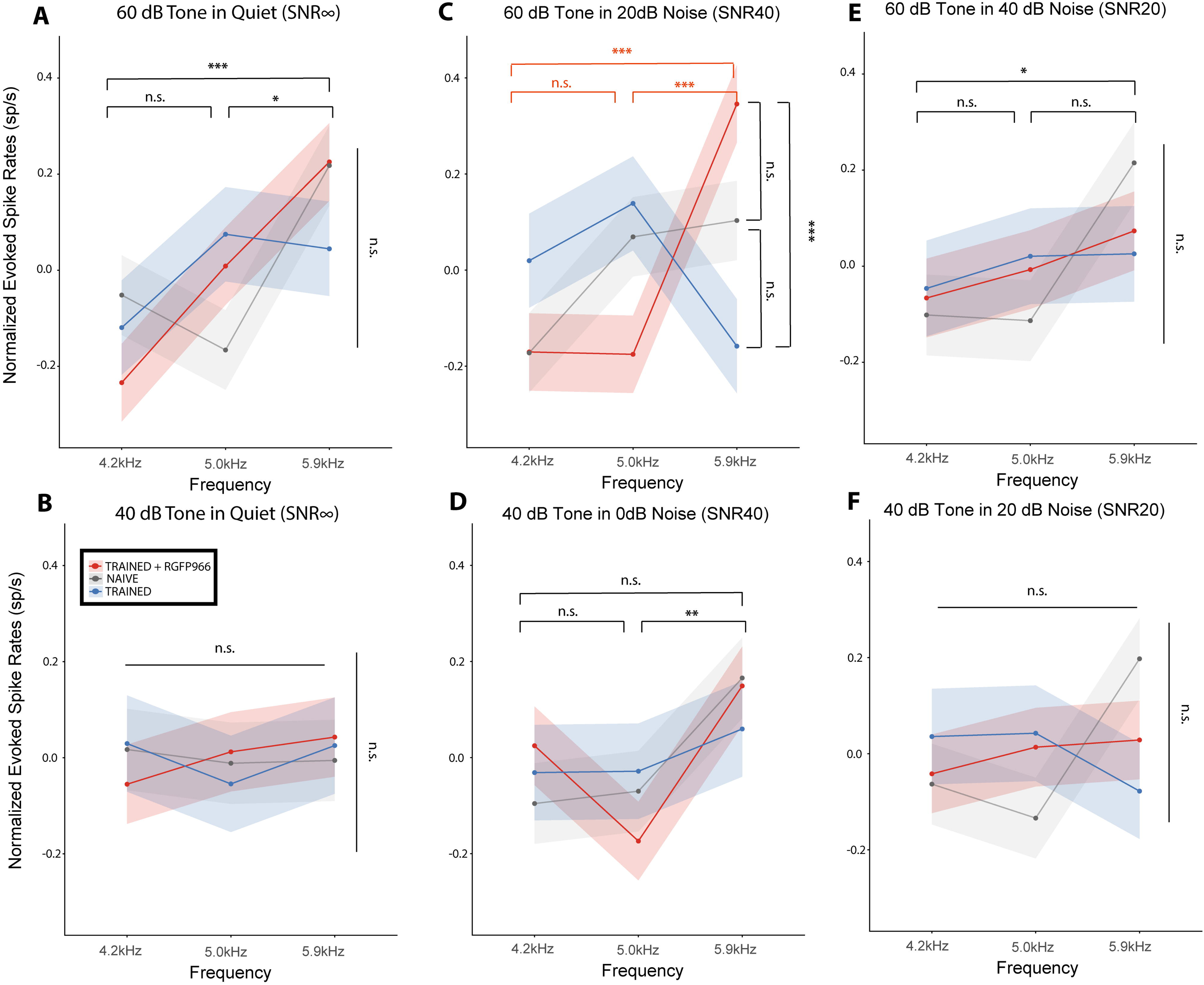
Learning- and noise-dependent modulation of auditory cortical frequency selectivity within the FRA region of interest. Frequency response areas (FRAs) were quantified for auditory cortical recording sites in trained rats (TRAINED and TRAINED+RGFP966) and in an untrained control group (NAÏVE; n = 7). Analyses were restricted to a predefined FRA region of interest (ROI; Fig. 2) comprising three test frequencies centered on the training tone (5.0 kHz) and adjacent novel frequencies (4.2 and 5.9 kHz), evaluated at the training sound level (60 dB SPL) and a lower sound level (40 dB SPL) across three background noise conditions (SNR∞, SNR40, SNR20). Spike rates were Z-scored across recording sites to normalize for between-site variability and enable group-level comparisons. Lines and shaded regions indicate group means SEM. **(A,B)** *Quiet (SNR*∞). At the training sound level (60 dB SPL; A), tone-evoked activity was frequency dependent across all groups, with the strongest responses observed at 5.9 kHz, independent of training or HDAC3 inhibition. At the lower sound level (40 dB SPL; B), frequency selectivity was abolished, and no effects of Group or Group × Frequency interactions were observed. **(C,D)** *Low background noise (SNR40).* Under low noise conditions at 60 dB SPL (C), a frequency-selective learning effect emerged that depended on HDAC3 inhibition. TRAINED+RGFP966 rats exhibited enhanced responses to the novel 5.9 kHz tone relative to both the training frequency (5.0 kHz) and the lower novel frequency (4.2 kHz), as well as relative to the untreated TRAINED group, reflected in a significant Frequency × Group interaction. This group-specific frequency selectivity was eliminated when tones were presented at 40 dB SPL (D), although responses remained strongest to 5.9 kHz across groups. **(E,F)** *High background noise (SNR20).* At the training sound level (60 dB SPL; E), frequency-dependent responding was weakened, and no effects of training or HDAC3 inhibition were detected despite a residual trend toward higher responses at 5.9 kHz. At the lower sound level (40 dB SPL; F), both frequency selectivity and group differences were fully abolished.

**Table 1.** Normalized Evoked Rate for Trained+RGFP966 and Trained animals. This table displays normalized evoked rate (M ± SE) for sites tuned to three test frequencies near the training tone frequency (i.e., 4.2kHz, 5.0kHz, and 5.9kHz) and at the training tone level (i.e., 60 dB SPL). Significant differences are bolded. *indicates a difference between vs. evoked by 4.2kHz, where *p<0.05, **p<0.01, and ***p<0.001. **§** indicates a difference vs. evoked by 5.0kHz, **^§^**p<0.05, **^§§^**p<0.01, and **^§§§^**p<0.001. **^¶^**indicates a difference vs. vehicle-treated animals, where **^¶^**p<0.05, **^¶¶^**p<0.01, and **^¶¶¶^**p<0.001.

| Frequency (kHz) | Group | Noise Condition |  |  |
| --- | --- | --- | --- | --- |
| | | Quiet ($SNR^\infty$) | Low Noise ($SNR_{40}$) | High Noise ($SNR_{20}$) |
| 4.2<br>60 dB | **Naïve** ($n = 7$ subjects; 94 sites) | $-0.05 \pm 0.083$ | $-0.17 \pm 0.082$ | $-0.10 \pm 0.084$ |
| | **Trained** ($n = 5$ subjects; 67 sites) | $-0.12 \pm 0.098$ | $0.02 \pm 0.097$ | $-0.04 \pm 0.099$ |
| | **Trained+RGFP966** ($n = 6$ subjects; 99 sites) | $-0.23 \pm 0.081$ | $-0.17 \pm 0.080$ | $-0.06 \pm 0.081$ |
| 5.0<br>60 dB | **Naïve** ($n = 7$ subjects; 94 sites) | $-0.16 \pm 0.083$ | $0.07 \pm 0.082$ | $-0.11 \pm 0.084$ |
| | **Trained** ($n = 5$ subjects; 67 sites) | $0.07 \pm 0.098$ | $0.14 \pm 0.097$ | $0.02 \pm 0.099$ |
| | **Trained+RGFP966** ($n = 6$ subjects; 99 sites) | $0.01 \pm 0.081$ | $-0.17 \pm 0.080$ | $-0.01 \pm 0.081$ |
| 5.9<br>60 dB | **Naïve** ($n = 7$ subjects; 94 sites) | $0.22 \pm 0.083$ | $0.10 \pm 0.082$ | $0.21 \pm 0.084$ |
| | **Trained** ($n = 5$ subjects; 67 sites) | $0.04 \pm 0.098$ | $-0.16 \pm 0.097$ | $0.02 \pm 0.099$ |
| | **Trained+RGFP966** ($n = 6$ subjects; 99 sites) | $0.22 \pm 0.081$ | **$0.34 \pm 0.080$** ***§§§¶¶¶ <sup>1</sup> | $0.07 \pm 0.081$ |

<sup>1</sup> Linear Mixed-Effects Model: Group: $F(2, 771) < 0.001$ , $p = 1.00$ ; Frequency: $F(2, 771) = 4.15$ , $p = 0.016$ ; Frequency x Group: $F(4, 771) = 6.55$ , $p < 0.001$ ; Tukey corrected t-tests for TRAINED+RGFP966: 5.0 kHz vs. 5.9 kHz: estimate = $-0.521$ , $t(514) = -4.57$ , $p < 0.0001$ §§§; 4.2 kHz vs. 5.9 kHz: estimate = $-0.516$ , $t(514) = -4.53$ , $p < 0.0001$ \*\*\*; for 5.9 kHz, TRAINED+RGFP966 vs. TRAINED group: Tukey corrected t-test: estimate = $0.504$ , $t(771) = 3.97$ , $p = 0.0002$ ¶¶¶

**Table 2.** Normalized Evoked Rate for Trained+RGFP966 and Trained animals. This table displays normalized evoked rate (M ± SE) for sites tuned to three test frequencies near the training tone frequency (i.e., 4.2kHz, 5.0kHz, and 5.9kHz) and near the training tone level (i.e., 40 dB SPL). Significant differences are bolded. *indicates a difference between vs. evoked by 4.2kHz, where *p<0.05, **p<0.01, and ***p<0.001. **§** indicates a difference vs. evoked by 5.0kHz, **^§^**p<0.05, **^§§^**p<0.01, and **^§§§^**p<0.001. **^¶^**indicates a difference vs. vehicle-treated animals, where **^¶^**p<0.05, **^¶¶^**p<0.01, and **^¶¶¶^**p<0.001.

| Frequency (kHz) | Group | Noise Condition |  |  |
| --- | --- | --- | --- | --- |
| | | Quiet ($SNR_{\infty}$) | Low Noise ($SNR_{40}$) | High Noise ($SNR_{20}$) |
| 4.2<br>40 dB | **Naïve** ($n = 7$ subjects; 94 sites) | $0.01 \pm 0.084$ | $-0.09 \pm 0.084$ | $-0.06 \pm 0.084$ |
| | **Trained** ($n = 5$ subjects; 67 sites) | $0.03 \pm 0.100$ | $-0.03 \pm 0.099$ | $0.03 \pm 0.099$ |
| | **Trained+RGFP966** ($n = 6$ subjects; 99 sites) | $-0.05 \pm 0.082$ | $0.02 \pm 0.081$ | $-0.04 \pm 0.082$ |
| 5.0<br>40 dB | **Naïve** ($n = 7$ subjects; 94 sites) | $-0.01 \pm 0.084$ | $-0.07 \pm 0.084$ | $-0.13 \pm 0.084$ |
| | **Trained** ($n = 5$ subjects; 67 sites) | $-0.05 \pm 0.100$ | $-0.02 \pm 0.099$ | $0.04 \pm 0.099$ |
| | **Trained+RGFP966** ($n = 6$ subjects; 99 sites) | $0.01 \pm 0.082$ | $-0.17 \pm 0.081$ | $0.01 \pm 0.082$ |
| 5.9<br>40 dB | **Naïve** ($n = 7$ subjects; 94 sites) | $-0.01 \pm 0.084$ | $0.16 \pm 0.084$ | $0.19 \pm 0.084$ |
| | **Trained** ($n = 5$ subjects; 67 sites) | $0.02 \pm 0.100$ | $0.059 \pm 0.099$ | $-0.07 \pm 0.099$ |
| | **Trained+RGFP966** ($n = 6$ subjects; 99 sites) | $0.04 \pm 0.082$ | $0.14 \pm 0.081$ | $0.02 \pm 0.082$ |

The effect of adding a novel background of low noise (SNR40) was investigated to contrast with tone-evoked activity in quiet conditions (SNR∞). Interestingly, a learning-induced effect on AC response bias emerged that was frequency selective under low levels of background noise in animals treated with the HDAC3 inhibitor during auditory learning (60 dB: Group: F(2, 771) < 0.001, p = 1.00; Frequency: F(2, 771) = 4.15, p = 0.016; Frequency x Group: F(4, 771) = 6.55, p < 0.001; Linear Mixed-Effects Model, **Fig. 3C**). Within TRAINED+RGFP966-treated rats, spike rates evoked by 5.9 kHz were higher compared to activity evoked both by 5.0 kHz and 4.2 kHz (Tukey corrected t-tests for TRAINED+RGFP966: 5.0 kHz vs. 5.9 kHz: estimate = -0.521, t(514) = -4.57, p < 0.0001; 4.2 kHz vs. 5.9 kHz: estimate = -0.516, t(514) = -4.53, p < 0.0001), which were also higher than tone-evoked activity to 5.9 kHz in the untreated TRAINED group (Tukey corrected t-test: estimate = 0.504, t(771) = 3.97, p = 0.0002; **Fig. 3C** and **Table 1**). Group differences in low background noise were abolished at lower sound levels of tone presentation (40 dB, Group: F(2, 771) < 0.001, p = 1.00; Frequency x Group: F(4, 771) = 0.81, p = 0.51; Mixed-Effects Model; **Fig. 3D**), though responding continued to be strongest to the 5.9 kHz tone (Frequency: F(2,771) = 4.75, p = 0.008; Tukey corrected posthoc: 5.0 kHz vs. 5.9 kHz: estimate = -0.215, t(514) = -2.97, p = 0.008; 4.2 kHz vs. 5.9 kHz: estimate = -0.159, t(514) = -2.19, p = 0.073; **Fig. 3D** and **Table 2**).

Strikingly, auditory cortical frequency selectivity in the TRAINED+RGFP966 group under low level background noise conditions was abolished in conditions of the highest levels of background noise (SNR20). Although there was a trend towards the same AC response bias to 5.9 kHz (60 dB, Frequency: F(2,771) = 3.26, p = 0.38; Tukey-corrected posthoc test: 4.2 kHz vs 5.9 kHz: estimate = -0.176, t(514) = -2.42, p = 0.04; 5.0 kHz vs 5.9 kHz: estimate = -0.138, t(514) = -1.902, p = 0.14; 4.2 kHz vs. 5.0 kHz: estimate = -0.038, t(514) = -0.52, p = 0.85), it did not depend on RGFP966 treatment or training (Group: F(2, 771) < 0.001, p = 1.00; Frequency x Group: F(4, 771) = 0.99, p = 0.41; Linear Mixed-Effects Model; **Fig. 3E** and **Table 1**). At lower sound levels of tone presentations in high noise, any evidence of group or frequency effects were eradicated (40 dBs, Frequency: F(2, 771) = 0.69, p = 0.49; Group: F(2, 771) < 0.001, p = 1.00; Frequency x Group: F(4, 771) = 1.93, p = 0.10; Linear Mixed-Effects Model; **Fig. 3F** and **Table 2**). Altogether, the data support that tone-reward learning induces a frequency-selective preservation of an auditory cortical response bias that can survive in low noise conditions and may be amplified under conditions of rapid task acquisition promoted by RGFP966 treatment.

### Auditory cortical responses to steady-state noise are suppressed in animals treated with an HDAC3 inhibitor during auditory learning

To explore the possibility that learning-induced effects mediated by RGFP966 could have effects on auditory cortical responses during steady-state noise alone, evoked activity during the time period *immediately prior* to tone onset was analyzed. Data are plotted across trials binned by the forthcoming tone frequency to-be-played on that trial for visualization. Overall, there was a suppression of auditory cortical responses to steady-state noise in trained animals when presented with 0 dB noise (Group: F(2,257) = 5.04, p = 0.007; **Fig. 4A**), with 20 dB noise (Group: F(2,257) = 5.18, p = 0.006; **Fig. 4B**), and with 40dB noise (Group: F(2,257) = 10.16, p < 0.001; **Fig. 4C**) that was most pronounced in TRAINED+RGFP966-treated animals (0 dB noise, Bonferroni corrected posthoc: TRAINED+RGFP966 vs. NAÏVE: t(191) = -3.17, p = 0.005; TRAINED+RGFP966 vs. TRAINED: t(164) = -1.35, p = 0.532; TRAINED vs. NAÏVE: t(159) = 1.52, p = 0.388; 20 dB noise, Bonferroni corrected posthoc: TRAINED+RGFP966 vs. NAÏVE: t(191) = -3.17, p = 0.005; TRAINED+RGFP966 vs. TRAINED: t(164) = -0.89, p = 1.00; TRAINED vs. NAÏVE: t(159) = 1.96, p = 0.151; 40 dB noise, Bonferroni corrected posthoc: TRAINED+RGFP966 vs. NAÏVE: t(191) = -4.50, p < 0.001; TRAINED+RGFP966 vs. TRAINED: t(164) = -2.11, p = 0.107; TRAINED vs. NAÏVE: t(159) = 1.97, p = 0.150; see **Table 3**). Together, these data suggest that a general outcome of auditory learning is to suppress auditory cortical responses to steady-state noise that can be strengthened under conditions of rapid task acquisition with RGFP966 treatment.

**Figure 4.**
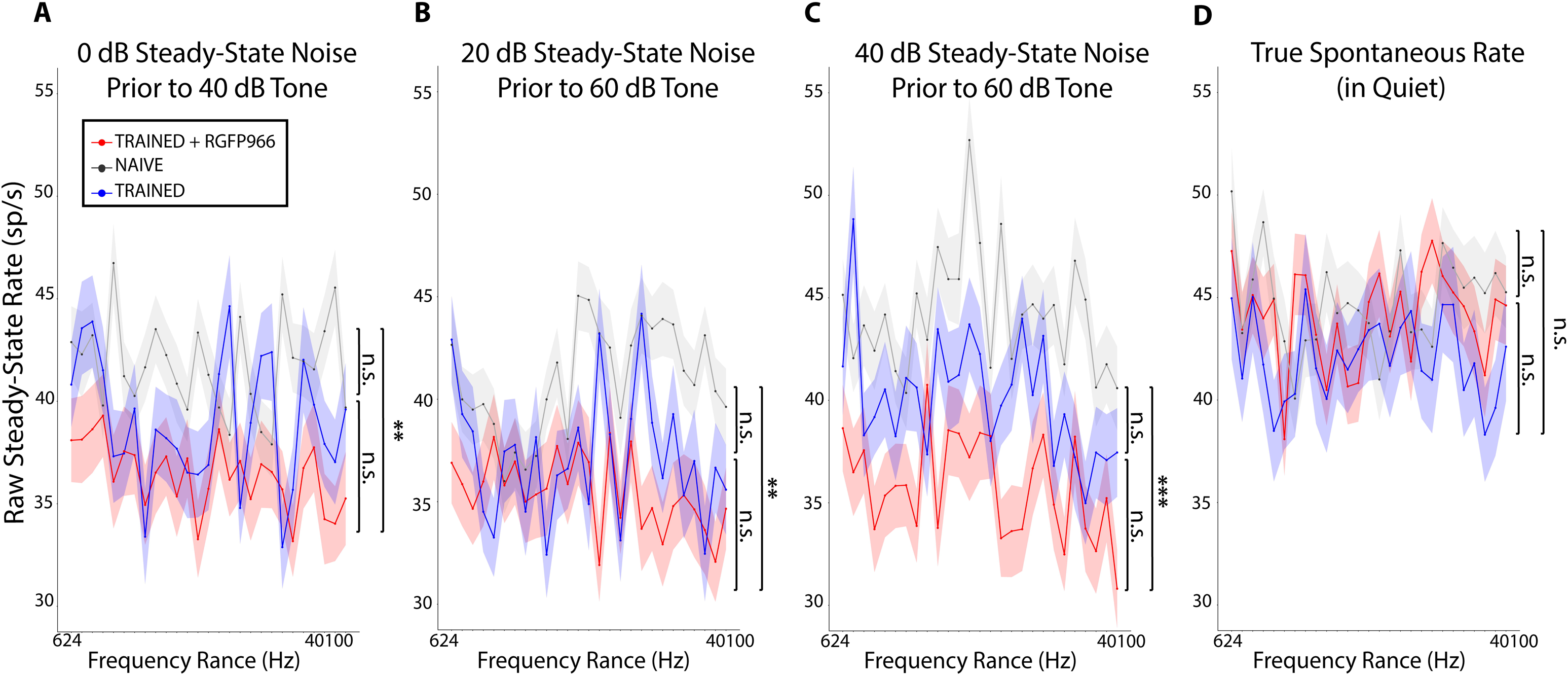
Auditory cortical responses to steady-state noise are suppressed by auditory learning and are enhanced by HDAC3 inhibition, without changes in spontaneous activity. Auditory cortical firing rates during steady-state background noise were quantified in trained rats (TRAINED and TRAINED+RGFP966) and untrained controls (NAÏVE). Analyses focused on the pre-tone epoch immediately preceding tone onset, isolating noise-evoked activity independent of tone-evoked responses. For visualization, data are plotted as a function of the forthcoming tone frequency on each trial. Lines and shaded regions denote group means ± SEM. (A–C) *Steady-state noise-evoked activity.* Mean firing rates during steady-state noise preceding tone presentation are shown for three noise levels: 0 dB noise prior to a 40 dB tone (A), 20 dB noise prior to a 60 dB tone (B), and 40 dB noise prior to a 60 dB tone (C). Across all three noise conditions, auditory cortical responses to steady-state noise were significantly reduced in trained animals, with the strongest suppression observed in TRAINED+RGFP966-treated rats. Group effects were present at all noise levels and were driven primarily by reduced activity in the TRAINED+RGFP966 group relative to NAÏVE animals, while differences between TRAINED and NAÏVE rats did not reach significance. Suppression of noise-evoked activity was progressively more pronounced as background noise level increased. (D) *Spontaneous activity in quiet.* True spontaneous firing rates measured during silent inter-trial intervals in quiet conditions did not differ between NAÏVE, TRAINED, and TRAINED+RGFP966 groups, indicating that training- and RGFP966-related reductions in activity are specific to sound-evoked responses during steady-state noise and do not reflect a global suppression of auditory cortical excitability.

**Table 3.** Steady-state noise and true spontaneous rate responses Trained+RGFP966 and Trained animals. The first three columns of this table display raw steady-state noise rate responses (M ± SE) in response to 0dB, 20 dB, and 40dB noise levels. The last column displays true spontaneous rate responses (M ± SE) in Quiet. Significant differences are bolded. *indicates a difference between vs. naïve animals where *p<0.05, **p<0.01, and ***p<0.001. **§** indicates a difference vs. trained animals, **^§^**p<0.05, **^§§^**p<0.01, and **^§§§^**p<0.001.

| Group | Noise Condition |  |  |  |
| --- | --- | --- | --- | --- |
|  | 0dB Noise | 20dB Noise | 40dB Noise | Quiet |
| **Naïve** ($n = 7$ subjects; 94 sites) | 41.8 ± 0.330 | 40.9 ± 0.329 | 44.2 ± 0.365 | 44.7 ± 0.368 |
| **Trained** ($n = 5$ subjects; 67 sites) | 38.9 ± 0.429 | 37.2 ± 0.425 | 40.1 ± 0.431 | 42.1 ± 0.477 |
| **Trained+RGFP966** ($n = 6$ subjects; 99 sites) | **36.4 ± 0.383\*\*<sup>2</sup>** | **35.5 ± 0.394\*\*<sup>3</sup>** | **35.8 ± 0.401\*\*\*<sup>4</sup>** | 44.0 ± 0.371 |

<sup>2</sup> One-way ANOVA: Group: $F(2,257)=5.04$ , $p=0.007$ ; Bonferroni corrected t-test: Trained+RGFP966 vs. Naïve: $t(191) = -3.17$ , $p = 0.005^{**}$ ; Trained+RGFP966 vs. Trained: $t(164) = -1.35$ , $p = 0.532$ ; Trained vs. Naïve: $t(159) = 1.52$ , $p = 0.388$ .
<sup>3</sup> One-way ANOVA: Group: $F(2,257)=5.18$ $p=0.006$ ; Bonferroni corrected t-test: Trained+RGFP966 vs. Naïve: $t(191) = -3.17$ , $p = 0.005^{**}$ ; Trained+RGFP966 vs. Trained: $t(164) = -0.89$ , $p = 1.00$ ; Trained vs. Naïve: $t(159) = 1.96$ , $p = 0.151$ .
<sup>4</sup> One-way ANOVA: Group: $F(2,257)=10.16$ , $p<0.001$ ; Bonferroni corrected t-test: Trained+RGFP966 vs. Naïve: $t(191) = -4.50$ , $p < 0.001^{***}$ ; Trained+RGFP966 vs. Trained: $t(164) = -2.11$ , $p = 0.107$ ; Trained vs. Naïve: $t(159) = 1.97$ , $p = 0.150$ .

### Spontaneous auditory cortical activity during quiet inter-stimulus intervals is not affected by training or treatment with an HDAC3 inhibitor

To investigate whether the suppressive effect on sound-evoked activity in trained and RGFP966-treated animals could be explained by an overall suppression of auditory cortical excitability, spontaneous auditory cortical activity during silent inter-trial intervals was analyzed. Spontaneous rate was not different between groups (F(2, 257) = 1.27, p = 0.282, **Fig. 4D** and **Table 3**). This finding bolsters the finding that lower cortical activity identified during periods of steady-state noise after training and RGFP966 treatment is most likely the result of a suppressive effect on noise-evoked auditory cortical activity *per se*.

### Background noise decreases behavioral task engagement

To evaluate how background noise may have had a general impact on behavioral engagement in the tone-reward task, a behavioral memory test was conducted to present tones under different SNRs that paralleled the background noise conditions used for electrophysiological recordings during auditory cortical mapping (see *Methods*). Bar-press responses to tones in both groups progressively decreased in graded steps between quiet and higher noise conditions (Noise Condition: F(4, 36) = 48.99, p < 0.001; Group: F(1, 9) = 0.13, p = 0.72; Noise Condition x Group: F(4, 36) = 0.41, p = 0.80, Mixed-ANOVA; **Fig. 5** and **Table 4**). Interestingly, the number of bar-presses that occurred in the highest noise condition (60 dB noise, SNR0) did not differ from the number of bar-presses made during trials of only white noise (i.e. 60 dB Gaussian noise) that did not include the presentation of a concurrent tone (Bonferroni corrected t-test: t(9) = -1.03, p = 1.00); both produced near-zero responding in the TRAINED+RGFP966 and TRAINED groups (see **Table 4)**. Because white noise and high noise (SNR0) conditions were statistically indistinguishable and yielded insufficient behavioral variance to estimate frequency selectivity, these conditions are reported descriptively but were excluded from mixed-effects analyses of frequency-selective memory in the next set of analyses.

**Figure 5.**
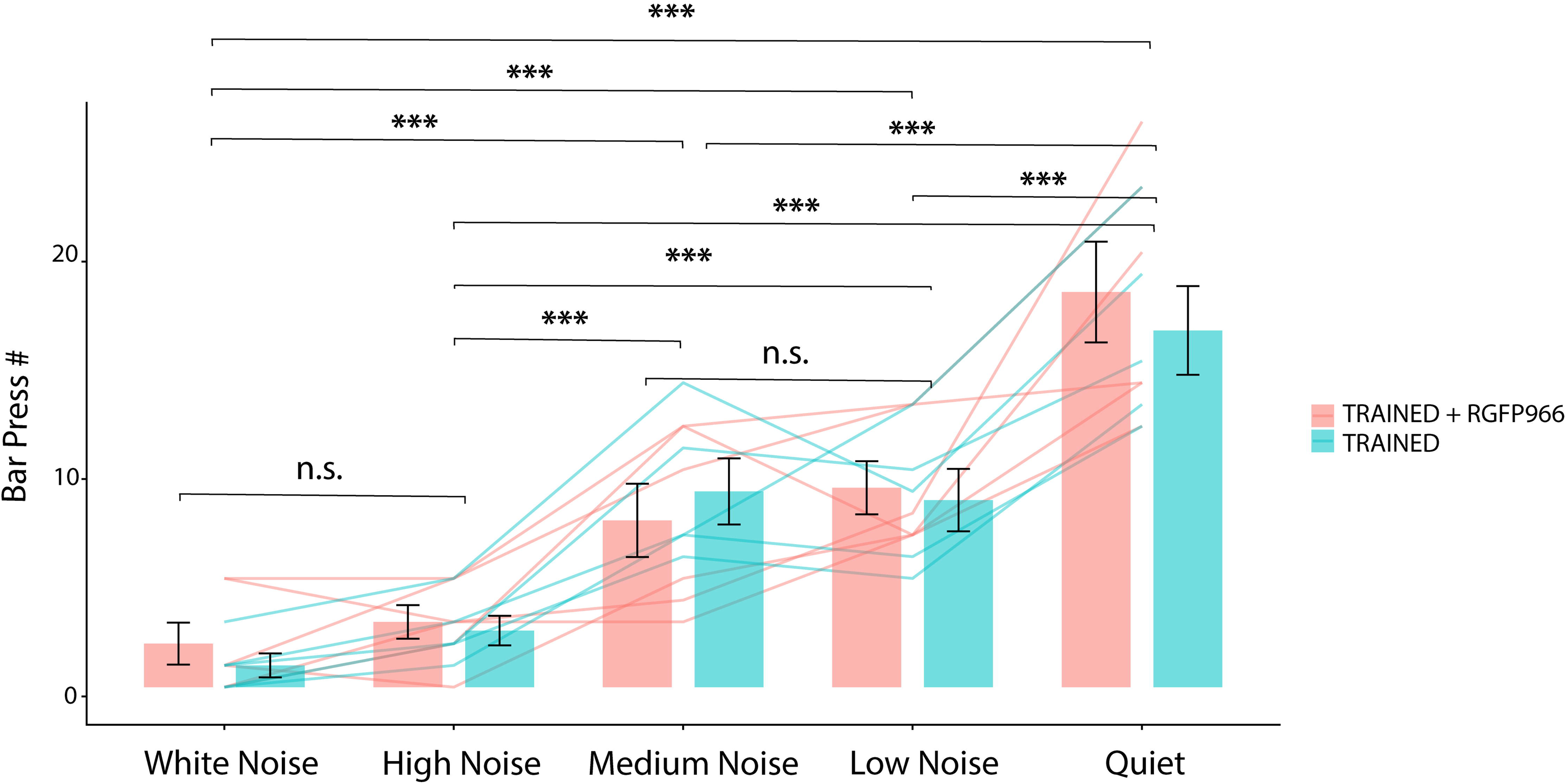
Background noise produces a graded reduction in behavioral responding during the tone-reward memory test, independent of HDAC3 inhibition. Rats previously trained on the tone-reward task (TRAINED and TRAINED+RGFP966) were tested with tone presentations embedded in progressively higher levels of background noise that paralleled the signal-to-noise ratio (SNR) conditions used during auditory cortical mapping (see *Methods*). Bars show group means ± SEM, with thin lines indicating individual animals. Because white noise and high-noise (SNR0) conditions were statistically indistinguishable and yielded minimal behavioral variance, they are shown for completeness but were excluded from subsequent mixed-effects analyses of frequency-selective memory performance.

**Table 4.** Raw number of bar presses during memory test for Trained+RGFP966 and Trained animals. This table displays raw number of bar presses (M ± SEM) in response to White Noise, High Noise, Medium Noise, Low Noise, and Quiet. *indicates a difference between noise conditions where *p<0.05, **p<0.01, and ***p<0.001.

| Group | Noise Condition |  |  |  |  |
| --- | --- | --- | --- | --- | --- |
|  | White Noise | High Noise | Medium Noise | Low Noise | Quiet |
| Trained<br>(n = 5 subjects) | 1.0 ± 0.55 | 2.6 ± 0.68 | 9.0 ± 1.52 | 8.6 ± 1.44 | 16.4 ± 2.04 |
| Trained+RGFP966<br>(n = 5 subjects) | 2.0 ± 0.09 | 3.0 ± 0.77 | 7.67 ± 1.69 | 9.17 ± 1.22 | 18.2 ± 2.32 |
| Comparisons | vs. HN: n.s.<sup>5</sup> | vs. MN: ***<sup>9</sup> | vs. LN: n.s.<sup>12</sup> | vs. Quiet: ***<sup>14</sup> |  |
|  | vs. MN: ***<sup>6</sup> | vs. LN: ***<sup>10</sup> | vs. Quiet: ***<sup>13</sup> |  |  |
|  | vs. LN: ***<sup>7</sup> | vs. Quiet: ***<sup>11</sup> |  |  |  |
|  | vs. Quiet: ***<sup>8</sup> |  |  |  |  |

<sup>5</sup> Mixed-ANOVA: Noise Condition x Group: F(4,36)=0.41, p=0.80; Group: F(1,9)=0.13, p=0.72; Noise Condition: F(4,36)=48.99, p<0.001; Bonferroni corrected t-test: White Noise vs. High Noise: t(9) = -1.03, p = 1.00 (n.s.)
<sup>6</sup> Mixed-ANOVA for all comparisons are as reported in (1); Bonferroni corrected t-test: White Noise vs. Medium Noise: t(9) = -5.41, p < 0.001\*\*\*
<sup>7</sup> Bonferroni corrected t-test: White Noise vs. Low Noise: t(9) = -5.85, p < 0.001\*\*\*
<sup>8</sup> Bonferroni corrected t-test: White Noise vs. Quiet: t(9) = -12.50, p < 0.001\*\*\*
<sup>9</sup> Bonferroni corrected t-test: High Noise vs. Medium Noise: t(9) = 4.38, p < 0.001\*\*\*
<sup>10</sup> Bonferroni corrected t-test: High Noise vs. Low Noise: t(9) = 4.82, p < 0.001\*\*\*
<sup>11</sup> Bonferroni corrected t-test: High Noise vs. Quiet Noise: t(9) = 11.47, p < 0.001\*\*\*
<sup>12</sup> Bonferroni corrected t-test: Medium Noise vs. Low Noise: t(9) = 0.43, p = 1.00
<sup>13</sup> Bonferroni corrected t-test: Medium Noise vs. Quiet Noise: t(9) = 7.09, p < 0.001\*\*\*
<sup>14</sup> Bonferroni corrected t-test: Low Noise vs. Quiet: t(9) = 6.65, p < 0.001\*\*\*

### The behavioral expression of frequency-specific memory recapitulates auditory cortical responses

A surprising finding in the auditory cortical datasets was the discovery that the 5.9 kHz test tone frequency evoked the highest rate of auditory cortical responding, including in low backgrounds of noise. To determine whether the selectivity of the cortical response to 5.9 kHz had an impact on behavioral response selectivity, the behavioral memory test conducted in the same animals was analyzed to determine the likelihood of responding to one of three different frequencies after achieving high levels of asymptotic performance. Given that the overall behavioral engagement was equivalent between treatment groups (as shown in **Fig. 5** and **Table 4)**, it was reasonable to use this analysis to approximate the effects of noise on frequency-selective tone detection in behavior. Memory testing included the training tone (5.0 kHz) and two nearby novel test frequencies (4.2 kHz and 5.9 kHz). Both TRAINED+RGFP966 and TRAINED groups displayed a frequency-selective behavioral response bias revealed by higher responding to the 5.9 kHz cue in quiet (SNR∞; Frequency: F(2,24) = 10.89, p = 0.0004; Group: F(1,24) < 0.001, p = 1.00; Group × Frequency: F(2,24) = 2.13, p = 0.14; Linear Mixed-Effects, **Fig. 6A** and **Table 5**). Frequency-selectivity for responding more to 5.9 kHz than to the other two tone frequencies was pronounced (planned contrasts averaged across groups: 5.9 kHz vs. 5.0 kHz, t(16) = 4.60, p = 0.0003; 5.9 kHz vs. the mean of 4.2 and 5.0 kHz, t(16) = 4.37, p = 0.0005). Interestingly, the addition of low-level background noise (20 dB noise; SNR40) paradoxically elevated the behavioral expression of frequency selectivity. Both groups showed markedly elevated responding to the 5.9 kHz test tone in both groups (Frequency: F(2, 24) = 15.20, p < 0.0001; Group: F(1, 24) < 0.0001, p = 1.00; Group × Frequency: F(2, 24) = 0.17, p = 0.85, Mixed-Effects Model; planned contrast comparing 5.9 kHz to the mean of 4.2 and 5.0 kHz, t(16) = 5.48, p = 0.0001; 5.9 kHz vs. 5.0 kHz, t(16) = 5.04, p = 0.0001; **Fig. 6B** and **Table 5**). In contrast, higher noise backgrounds (40 dB noise; SNR20) disrupted the expression of signal-specific memory toward the 5.9-kHz cue. Neither tone frequency nor treatment significantly influenced responding (Frequency: F(2,24) = 1.85, p = 0.18; Group: F(1, 24) < 0.001, p = 1.00; Group × Frequency: F(2, 24) = 0.09, p = 0.92; **Fig. 6C** and **Table 5**). In line with the prior reported finding of poor behavioral engagement during high noise conditions, frequency-selective contrasts collapsed across groups did not reach significance (t(16) = 1.68, p = 0.11), indicating that the highest noise tested condition completed abolished the expression of behavioral response selectivity for tone frequency that had emerged under low noise conditions. The behavioral response bias to the 5.9 kHz tone is a remarkable match to the auditory cortical tone-evoked response patterns. This match is even more remarkable when considering that the sound frequency presented during tone-reward task training was 5.0 kHz, while the behavioral and neural response bias were both peak-shifted to the 5.9 kHz frequency. Even more remarkable was the match observed in auditory cortical and behavioral nonlinearity to enhance and then abolish frequency-selective response biases with increasing background noise conditions from low to high, respectively.

**Figure 6.**
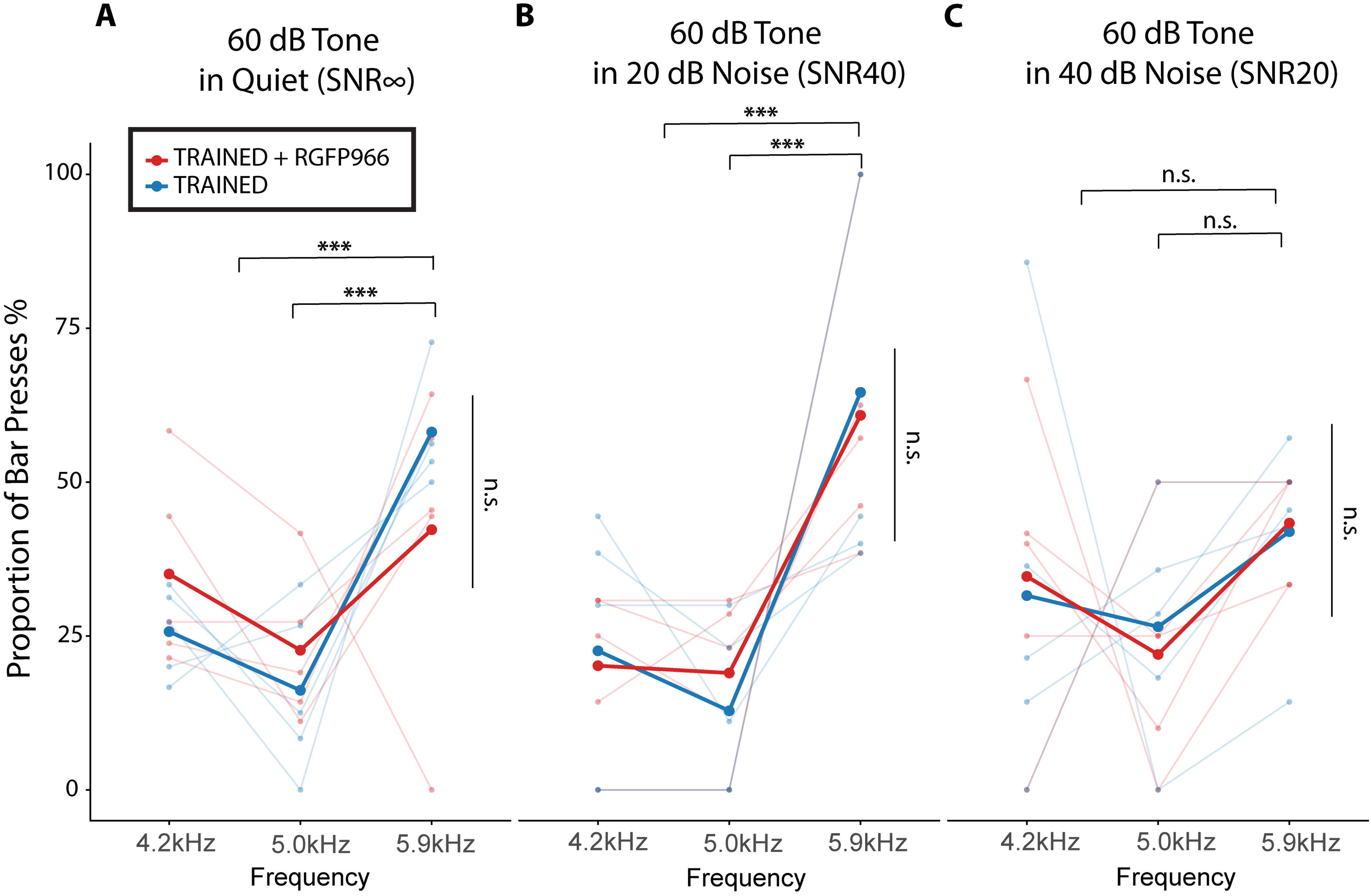
Behavioral expression of frequency-specific memory mirrors auditory cortical response selectivity and its modulation by background noise. Memory tests were conducted in the same animals used for physiological analyses and included the training frequency (5.0 kHz) and two nearby novel frequencies (4.2 and 5.9 kHz). Proportion of bar-press responses is shown for TRAINED and TRAINED+RGFP966 groups; thin lines indicate individual animals, and thick lines denote group means. (A) *quiet (SNR*∞). In the absence of background noise, both groups exhibited a frequency-selective behavioral response bias, with significantly higher responding to the 5.9 kHz tone compared with the training frequency (5.0 kHz) and the mean of training frequency and lower novel frequency (4.2 kHz). This effect was independent of RGFP966 treatment, as evidenced by a significant main effect of Frequency without Group or Group × Frequency interactions. (B) *Low background noise (SNR40; 20 dB noise).* The addition of low-level background noise paradoxically enhanced the behavioral expression of frequency selectivity. Both TRAINED and TRAINED+RGFP966 animals showed markedly elevated responding to the 5.9 kHz tone relative to the other frequencies, yielding a stronger main effect of Frequency while remaining independent of treatment group. (C) *Higher background noise (SNR20; 40 dB noise).* In contrast, higher levels of background noise abolished frequency-selective responding.

**Table 5.** Proportion of bar-press responses for Trained+RGFP966 and Trained animals. This table displays proportion of bar-press responses (M ± SEM) at three test frequencies near the training tone frequency (i.e., 4.2kHz, 5.0kHz, and 5.9kHz) and at the training tone level (i.e., 60 dB SPL). *indicates a significant effect where *p<0.05, **p<0.01, and ***p<0.001.

| Frequency (kHz) | Group | Noise Condition |  |  |
| --- | --- | --- | --- | --- |
| | | Quiet ($SNR_{\infty}$) | Low Noise ($SNR_{40}$) | Medium Noise ($SNR_{20}$) |
| 4.2 kHz<br>60 dB | **Trained**<br>($n = 5$ subjects) | 25.7 ± 3.21 | 22.6 ± 9.50 | 31.6 ± 14.8 |
| | **Trained+RGFP966**<br>($n = 5$ subjects) | 35.1 ± 7.08 | 20.2 ± 5.87 | 34.7 ± 10.9 |
| 5.0 kHz<br>60 dB | **Trained**<br>($n = 5$ subjects) | 16.2 ± 6.09 | 12.8 ± 6.05 | 26.5 ± 8.41 |
| | **Trained+RGFP966**<br>($n = 5$ subjects) | 22.7 ± 5.47 | 19.0 ± 5.70 | 22.0 ± 8.46 |
| 5.9 kHz<br>60 dB | **Trained**<br>($n = 5$ subjects) | 58.1 ± 3.91 | 64.6 ± 14.5 | 41.9 ± 7.33 |
| | **Trained+RGFP966**<br>($n = 5$ subjects) | 42.3 ± 11.2 | 60.9 ± 10.06 | 43.3 ± 4.08 |
| *Linear Mixed-Effects Model:* | | Frequency:<br>$F(2,24) = 10.89$,<br>**$p = 0.0004^{***}$** | Frequency:<br>$F(2, 24) = 15.20$,<br>**$p < 0.0001^{***}$** | Frequency:<br>$F(2,24) = 1.85$,<br>$p = 0.18$ |
| | | Group:<br>$F(1,24) < 0.001$,<br>$p = 1.00$ | Group:<br>$F(1, 24) < 0.0001$,<br>$p = 1.00$ | Group:<br>$F(1, 24) < 0.001$,<br>$p = 1.00$ |
| | | Group × Frequency:<br>$F(2,24) = 2.13$, $p = 0.14$ | Group × Frequency:<br>$F(2,24) = 0.17$, $p = 0.85$ | Group × Frequency:<br>$F(2,24) = 0.09$, $p = 0.92$ |
| *Planned contrast posthocs:* | | 5.9 kHz vs. 5.0 kHz: $t(16) = 4.60$,<br>**$p = 0.0003^{***}$** | 5.9 kHz vs. 5.0 kHz: $t(16) = 5.04$,<br>**$p = 0.0001^{***}$** | |
| | | 5.9 kHz vs. mean of 4.2 & 5.0 kHz: $t(16) = 4.37$,<br>**$p = 0.0005^{***}$** | 5.9 kHz vs. mean of 4.2 & 5.0 kHz: $t(16) = 5.48$,<br>**$p = 0.0001^{***}$** | |

## DISCUSSION

The present study identifies learning-dependent mechanisms that shape how auditory cortex encodes behaviorally relevant signals. Across behavioral and electrophysiological measures, background noise degraded expression of a learned sound-specific memory, consistent with extensive human literature showing that speech and word recognition deteriorate as noise levels increase despite constant SNR <u>(e.g., Pollack and Pickett, 1958; Dubno et al., 2005)</u>. The findings show that associative learning reorganizes frequency-specific cortical representations in ways that remain detectable—and in some cases amplified—under modest acoustic interference. Pharmacological inhibition of HDAC3 strengthens this representational plasticity by selectively enhancing AC responses evoked by the learned signal while suppressing noise-driven responsivity. Together, the results reveal a coordinated pattern of plasticity in which associative learning and HDAC3-regulated neuroplasticity interact to improve cortical signal processing and behavior under degraded acoustic conditions. These findings extend classical views of AC plasticity by demonstrating that expression of learned representations is not fixed, but instead emerges through dynamic interactions between prior learning and ongoing acoustic context (Brandon et al., 2003). More broadly, HDAC3-regulated neuroplasticity may determine when and under what sensory conditions cortical plasticity can support sound-guided behavior.

A central result is that frequency-selective AC response biases were functionally latent under quiet conditions and became robustly expressed under low levels of background noise, particularly in RGFP966-treated learners. This demonstrates that noise can serve as a probe for latent learning-dependent tuning changes, analogous to latent learning frameworks proposed in other systems (Lubow and Moore, 1959; Kuchibhotla et al., 2019; Drieu et al., 2025). Under low noise (SNR40), the response bias toward 5.9 kHz was amplified, whereas under higher noise (SNR20) this selectivity collapsed. This nonlinear pattern is consistent with the idea that background noise interacts with the stability of learned cortical representations, revealing a threshold beyond which noise disrupts expression of plasticity-driven memory traces. These findings support a model in which learning-induced frequency-selective plasticity confers robustness only within a constrained dynamic noise range, beyond which cortical suppression mechanisms mask frequency-selective coding. Thus, learned auditory memories are not expressed uniformly across contexts but depend on the interplay between representational strength and sensory context including interference. An interaction between noise-driven and RGFP966-driven changes in AC coding further reveals a mechanistic convergence. Prior work has shown that background noise can paradoxically improve discrimination between adjacent frequencies by reducing receptive field overlap (Christensen et al., 2019), while treatment with HDAC3 inhibition during associative learning similarly narrows signal-specific tuning bandwidth (Bieszczad et al., 2015; Shang et al., 2019; Rotondo and Bieszczad, 2020, 2021a). In the present study, HDAC3 inhibition preserved and enhanced learning-induced frequency selectivity for the learned tone amidst background noise. Together, the findings suggest that HDAC3-mediated regulation stabilizes key coding features like tuning selectivity and enhanced sound-evoked responses that benefit signal-in-noise detection.

A second major finding is that HDAC3 inhibition selectively suppressed steady-state noise responses without altering spontaneous cortical activity. This indicates that HDAC3-dependent plasticity enhances cortical signal-to-noise ratios not by increasing excitability broadly, but by shaping how AC responds to noise. Across noise levels, RGFP966-treated animals showed reduced noise-evoked firing relative to naïve animals and untreated learners, even though noise was absent during acquisition. The lack of spontaneous firing differences argues against global hyperexcitability as a mechanism (Christensen et al., 2025) and instead supports a shift in excitatory–inhibitory balance that prioritizes responses to remembered signals while suppressing non-informative background noise (Teschner et al., 2016; Christensen et al., 2019; Lakunina et al., 2022; Mischler et al., 2023). This redistribution of cortical responsivity prioritizes behaviorally relevant information while minimizing effort costs associated with noise processing.

An intriguing implication concerns whether animals with *a priori* cortical properties, such as reduced noise responsivity, would facilitate faster acquisition. While RGFP966 treatment correlated with accelerated learning across the cohort, it remains possible that animals predisposed to low noise-driven cortical excitability experience an inherently improved neural signal-to-noise ratio during learning. More broadly, these findings raise the hypothesis that mitigating noise during learning via biological or environmental manipulations could facilitate learning and memory outcomes. This hypothesis has support in broadly relevant educational and clinical settings (Bronzaft, 1981; Alain et al., 2014; Kuchinsky et al., 2016; Krizman et al., 2017; Hennessy et al., 2021), suggesting that reducing acoustic interference may disproportionately benefit individuals with inherently weaker cortical suppression mechanisms.

The enhanced expression of frequency-selective biases under low noise in RGFP966-treated animals further highlights that HDAC3-regulated plasticity stabilizes high-fidelity representations that remain decodable under moderately degraded conditions. However, it does not render cortical processing noise-proof. At higher noise levels, frequency selectivity was abolished in treated and untreated animals alike, indicating that HDAC3-regulated plasticity extends, but does not remove, the operational limits under which learned auditory representations can be expressed. This distinction is mechanistically important: HDAC3 inhibition increases representational robustness without overriding physiological constraints imposed by strong sensory interference. Learning may lessen a sound’s vulnerability to noise by enhancing memory trace stability, potentially promoting greater separation between neural populations responding to learned signals, novel sounds, and background noise. For example, stream segregation depends critically on prior knowledge, and memory for a sound can enhance its segregation from background noise (Bregman, 1994; Snyder and Alain, 2007). This interpretation aligns with evidence that sparse neural populations in auditory cortex can encode behaviorally relevant sounds in a noise-invariant manner up to a limit (Schneider and Woolley, 2013; Kang and Kanold, 2023). The present study extends this literature by linking learning experience and behavioral outcomes directly to AC physiology under noisy conditions, supporting a model in which HDAC3 regulates learning-induced representational strength and the operational limits of functional expression.

A consistent observation across neural and behavioral datasets was a peak-shifted responsivity bias toward the 5.9 kHz test frequency rather than the 5.0 kHz training frequency. This shift reflects a canonical signature of associative generalization and receptive field reweighting, consistent with classical behavioral accounts of peak shift (Spence, 1937; <u>Purtle, 1973)</u> and with evidence that auditory learning reshapes population coding around behaviorally salient rather than veridically trained features (Wisniewski et al., 2009; Miasnikov and Weinberger, 2012). The convergence of cortical and behavioral biases toward 5.9 kHz indicates that learning modified representational structure in auditory cortex to prioritize discriminability over stimulus accuracy. Such representational biases are thought to emerge from transient neural states engaged during learning that shape how sensory information is encoded and later expressed. Stimulus-evoked gamma activity during learning predicts later behavioral responding to both reinforced and nearby non-reinforced frequencies, suggesting that encoding-related cortical dynamics can bias internally generated or inferred representations (Gonsalves and Paller, 2000; Headley and Weinberger, 2011). Within this framework, peak shift may arise when encoding processes favor expected or behaviorally relevant features over exact acoustic identity. HDAC3-regulated plasticity may stabilize the impact of these encoding-related processes, extending the conditions under which such biased representations remain decodable in noise. The SNR-dependent modulation of peak shift further reinforces that learned representational bias is dynamically gated by sensory context.

HDAC3 inhibition thus emerges as a mechanistic probe for dissecting the cortical computations underlying listening in noise. By enhancing consolidation of frequency-specific learning without globally increasing excitability, RGFP966 reveals candidate neural features sufficient for signal detection under challenging conditions. These findings support the view that HDAC3 functions as a molecular gatekeeper regulating both the speed at which learning influences cortical computation and the stability with which those changes are expressed under increasing sensory interference. Future work should determine whether HDAC3-dependent preservation of AC coding predicts sound-specific behavior in noise within individuals, and whether similar mechanisms extend to temporal coding features relevant for remembering complex acoustic signals (Phan et al., 2017; Rotondo and Bieszczad, 2021b). Extending this work to aged animals will also be critical to assess whether HDAC3-mediated robustness can mitigate age-related decline in listening abilities (Kuchinsky et al., 2016; Burns and Gräff, 2021; Shilling-Scrivo et al., 2021).

Overall, these findings support an integrated mechanistic model in which auditory learning shapes cortical processing to support listening in noise. Learning produces frequency-specific plasticity that enhances signal detection under modest interference while engaging complementary processes that suppress steady-state noise. HDAC3 inhibition strengthens these learning-induced processes and extends the range of acoustic conditions over which memory can be expressed. By linking epigenetic mechanisms to cortical coding dynamics and sound-guided behavior, this study broadens our understanding of how plasticity is stabilized and deployed across sensory contexts. Molecular constraints on learning and memory consolidation may influence not only the persistence of cue memories (Campbell and Wood, 2019; Shang and Bieszczad, 2022) but also their expression in real-world environments.

## Contributions

KMB & NA designed research; NA performed research, KMB & NA contributed unpublished analytic tools; NA analyzed data; KMB & NA wrote the paper.

## Acknowledgements

This work was supported by the National Institutes of Health, the National Institute on Deafness and Communication Disorders [R01-DC018561 to K.M.B.] and funding from the School of Arts and Sciences at Rutgers University—New Brunswick. The authors wish to thank Dr. Justin Yao for helpful discussions that motivated analyses that contributed to the completion of this work, and Dr. Mimi Phan for reviewing an initial version of the manuscript. ChatGPT 5.2 was used for editing purposes to adhere to word count limits in the abstract, introduction and discussion sections of this manuscript. We thank former and current members of the CLEF Lab, especially Dr. Andrea Shang, who generated initial versions of the analytical code that was adapted for investigating auditory cortical response profiles across noise conditions; and to Rebecca Shear, who contributed to animal training across the large cohort reported, and to Elias Youssef, M.S., who offered technical assistance throughout the project. The authors are also grateful to the membership of Rutgers Auditory Neuroscience Research Initiative (RANRI) for their continuous insight and valuable input to this project.

## SUPPLEMENTARY FIGURE LEGENDS

**Figure S1.**
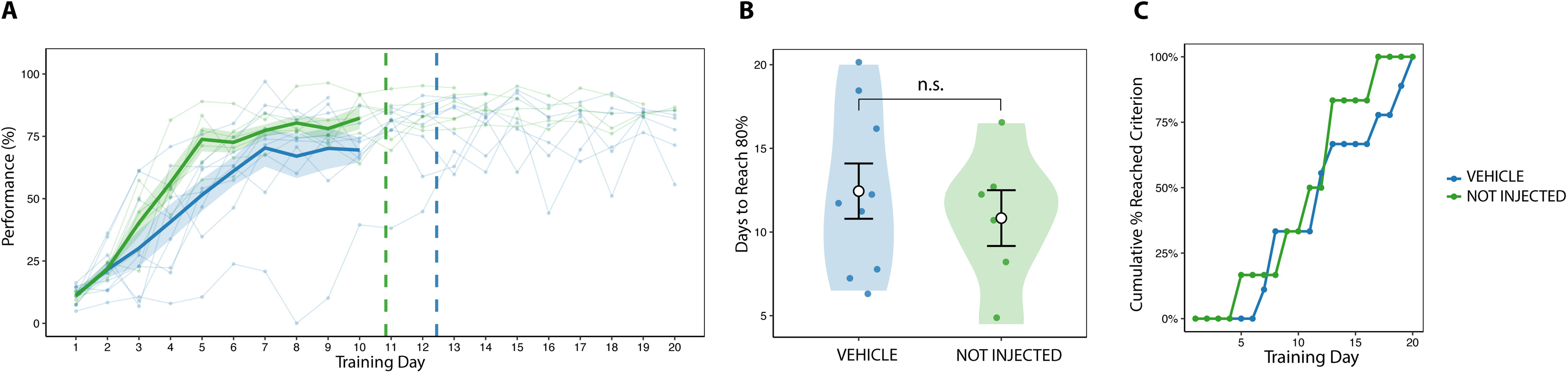
Vehicle injections do not alter acquisition of the tone-reward task. **(A)** Acquisition curves showing daily performance across training days for vehicle-injected and non-injected trained animals. Thin lines represent individual animals, and thick lines indicate group means ± SEM. Dashed vertical lines denote the mean day at which each group reached the 80% performance criterion. Both groups exhibited similar learning trajectories and reached asymptotic performance over comparable time courses. **(B)** Days required to reach the 80% performance criterion for vehicle-injected and non-injected animals. Individual animals are shown with group means ± SEM overlaid; no significant difference was observed between groups. **(C)** Cumulative percentage of animals reaching the 80% criterion as a function of training day, illustrating overlapping acquisition dynamics for vehicle-injected and non-injected animals. The mean number of days to reach criterion did not differ between vehicle-injected and non-injected rats (Wilcoxon rank-sum test, p = 0.72).

**Figure S2.**
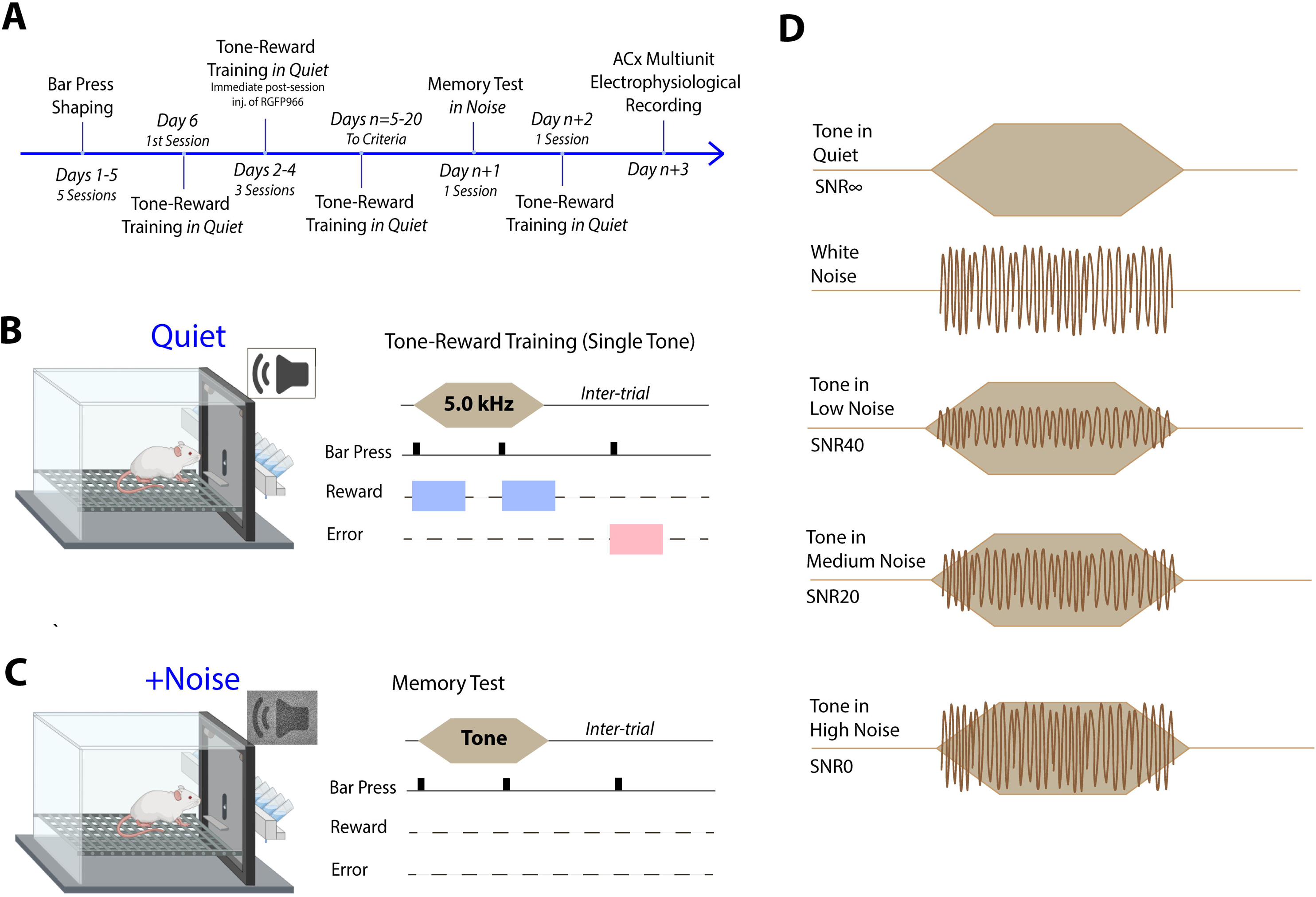
Experimental timeline for learning the tone-reward task, memory testing, and auditory cortical electrophysiological recording. (A) *Experimental timeline.* Animals underwent bar-press shaping followed by tone-reward training in quiet. Post-session injections of the selective HDAC3 inhibitor RGFP966 or vehicle treatment were administered immediately after training sessions on days 2, 3 and 4. Training in quiet conditions continued daily until animals reached the acquisition criterion. Upon attaining criterion, animals completed a behavioral memory test in a novel background of different noise conditions, followed 24-48 hrs later by a final reinforced training session in quiet to confirm stable task performance prior to auditory cortical electrophysiological recordings. (B) *Tone-reward training in quiet.* During training sessions, a single 5.0 kHz tone was presented in quiet. Bar presses occurring during tone presentation were reinforced with water reward, whereas responses outside the tone window were not reinforced. (C) *Memory test in noise.* To ensure stable task engagement immediately prior to memory testing, animals completed a brief reinforced warm-up session (∼5 min) that was immediately followed by the memory test. During the memory test, tones were presented without reinforcement, and bar-press responses were recorded as a measure of memory-guided behavior. (D) *Acoustic structure of memory test stimuli.* Memory test trials consisted of an 8-s tone presented either in quiet (SNR∞) or embedded in notched Gaussian white noise at three signal-to-noise ratios: low noise (60 dB tone + 20 dB noise; SNR40), medium noise (60 dB tone + 40 dB noise; SNR20), and high noise (60 dB tone + 60 dB noise; SNR0).

**Figure S3.**
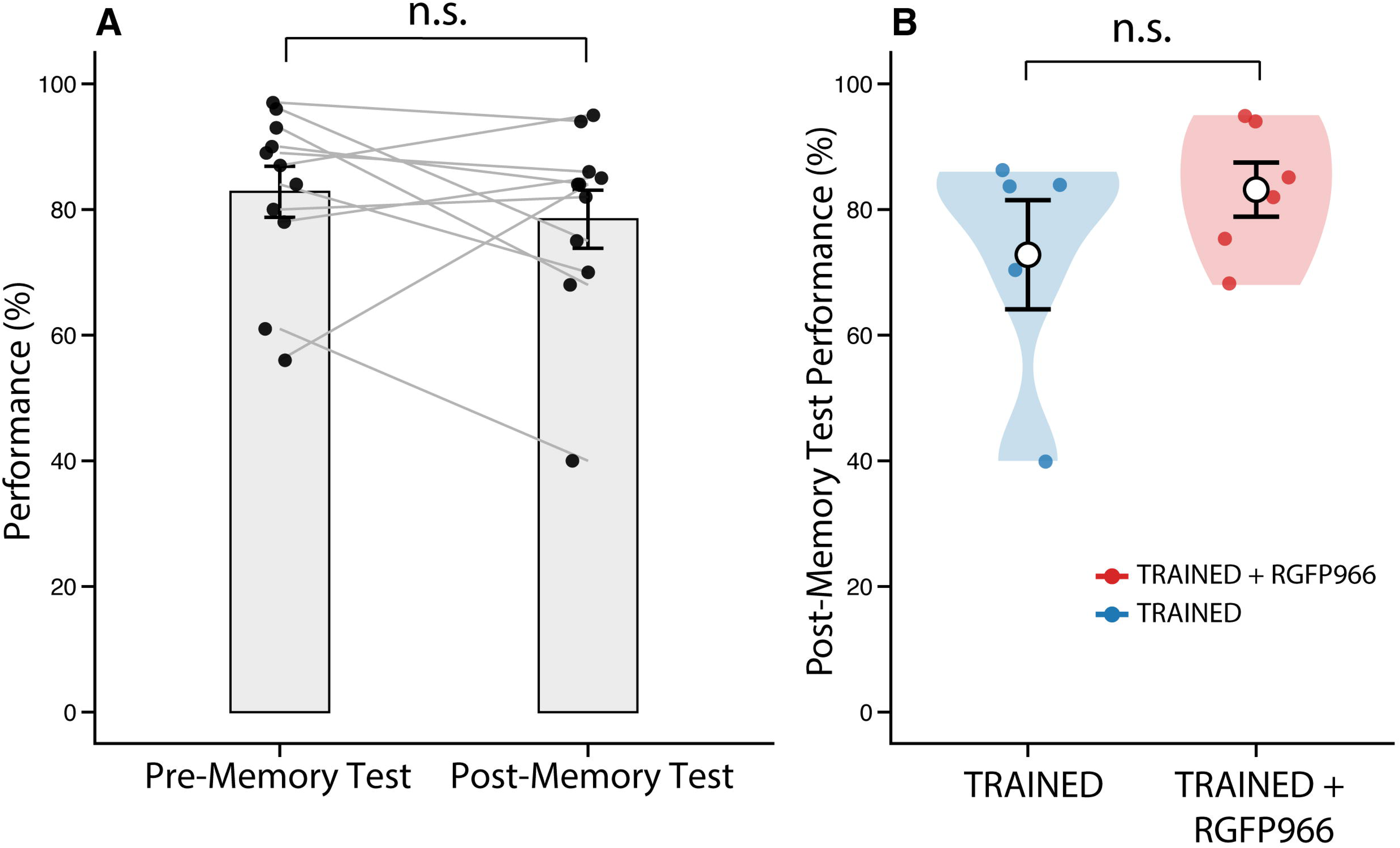
Memory testing does not disrupt subsequent tone–reward task performance. All animals completed an additional tone-reward training session 24-48 hrs after memory testing. **(A)** *Within-animal performance stability.* Performance during the final reinforced training trial conducted before memory testing was compared with performance during the reinforced session conducted after memory testing. Individual animals are shown with paired data points connected by lines; bars represent group means ± SEM. Performance levels did not differ between pre- and post-memory test sessions, indicating that memory test did not disrupt task performance. **(B)** *Between-group comparison after memory testing.* Performance during the post-memory test training session is shown separately for TRAINED and TRAINED+RGFP966 animals. Individual animals are overlaid on violin plots depicting the distribution of performance values, with group means ± SEM indicated. Final performance did not differ between groups, demonstrating equivalent task proficiency following memory testing.

